# Maternal genetic affinities of Koṅkaṇī population in the southwest coast of India

**DOI:** 10.1101/2024.09.23.614647

**Authors:** Jaison Jeevan Sequeira, Lomous Kumar, George van Driem, Kumarasamy Thangaraj, Mohammed S Mustak

## Abstract

Koṅkaṇ region on the west coast of India is a hotspot of culture, folklore and ethnolinguistic diversity. The genetic landscape of this region remains understudied. The present study features Koṅkaṇī population residing along the Koṅkaṇ Malabar coast. We have sequenced complete mitogenomes of 85 and the hypervariable region of 210 Koṅkaṇī individuals to understand the maternal gene pool of this region. Comparative analysis of the over 5000 mitogenomes revealed that the Koṅkaṇī population clustered at a convergence point on the PCA plot, presumably due to a diverse maternal gene pool with both autochthonous and West Eurasian components. A distinct clustering pattern was observed within the subgroups of Sārasvata and non-Sārasvata Koṅkaṇī groups, indicating unique ancestral maternal lineages in them. This distinction is majorly due to the N macrohaplogroup lineages found in this population. We observe low haplotype and nucleotide diversity in Citrapur Sārasvata Brahmins (CSB), Rājāpur Sārasvata Brahmins (RSB), Khārvi and Kuḍubi compared to Gauḍa Sārasvata Brahmins (GSB) and Roman Catholics. The assimilation of both pre and post Last Glacial Maximum (LGM) haplogroups like M57, M36, M37, M3, M30, R8 and U2 in the Koṅkaṇī population suggests active movement and settlement along the Koṅkaṇ region on the west coast of India since the Late Pleistocene through the Holocene.

## 1. Introduction

India is a hotspot of genomic diversity and home to peoples of various castes, ethnicities, religious communities and linguistic groups, including major language families such as Indo-European, Dravidian, Austroasiatic and Trans-Himalayan. The phylogeographies of genes and language families are correlated (Cavalli-Sforza et al. 1995), and this correlation is observed most neatly with paternally inherited markers, and much less so with mitochondrial or autosomal markers, with a number of notable exceptions (Poloni et al. 1997; Belledi et al. 2000; van Driem 2021).

On the southwestern coast of India lies the Koṅkaṇ region, including all the land between the Western Ghats and the Indian Ocean, from the latitude of Daman in the north to the Gaṅgavalli river and Carwar (*Kārvār*) in the south. Koṅkaṇī language is spoken all along this narrow strip of land with considerable local differences in phonetics, grammar and vocabulary. Outside of this main Koṅkaṇī speaking stretch of coast, the language is also spoken by scattered Koṅkaṇī communities in the littoral region, extending up to Mālvaṇ in the north and down to Mangalore in the south, including parts of Kerala and Bombay (Fig. 1a). According to the 2011 census data, there are nearly 22.5 lakh Koṅkaṇī speakers in India. Conventionally, the Koṅkaṇī language has been viewed as a member of the southwestern branch of Indo-Aryan, which furthermore comprises Marāṭhī, Gujarātī, Divehi and Sinhalese, whereby Koṅkaṇī, both geographically and linguistically, occupies an intermediate position between Marāṭhī and Gujarātī on the Indian mainland and Divehi and Sinhalese on the islands. The Koṅkaṇī linguistic area itself can be divided into northern and southern dialect areas, in addition to a few community-specific Koṅkaṇī sociolects.

**Fig. 1.**
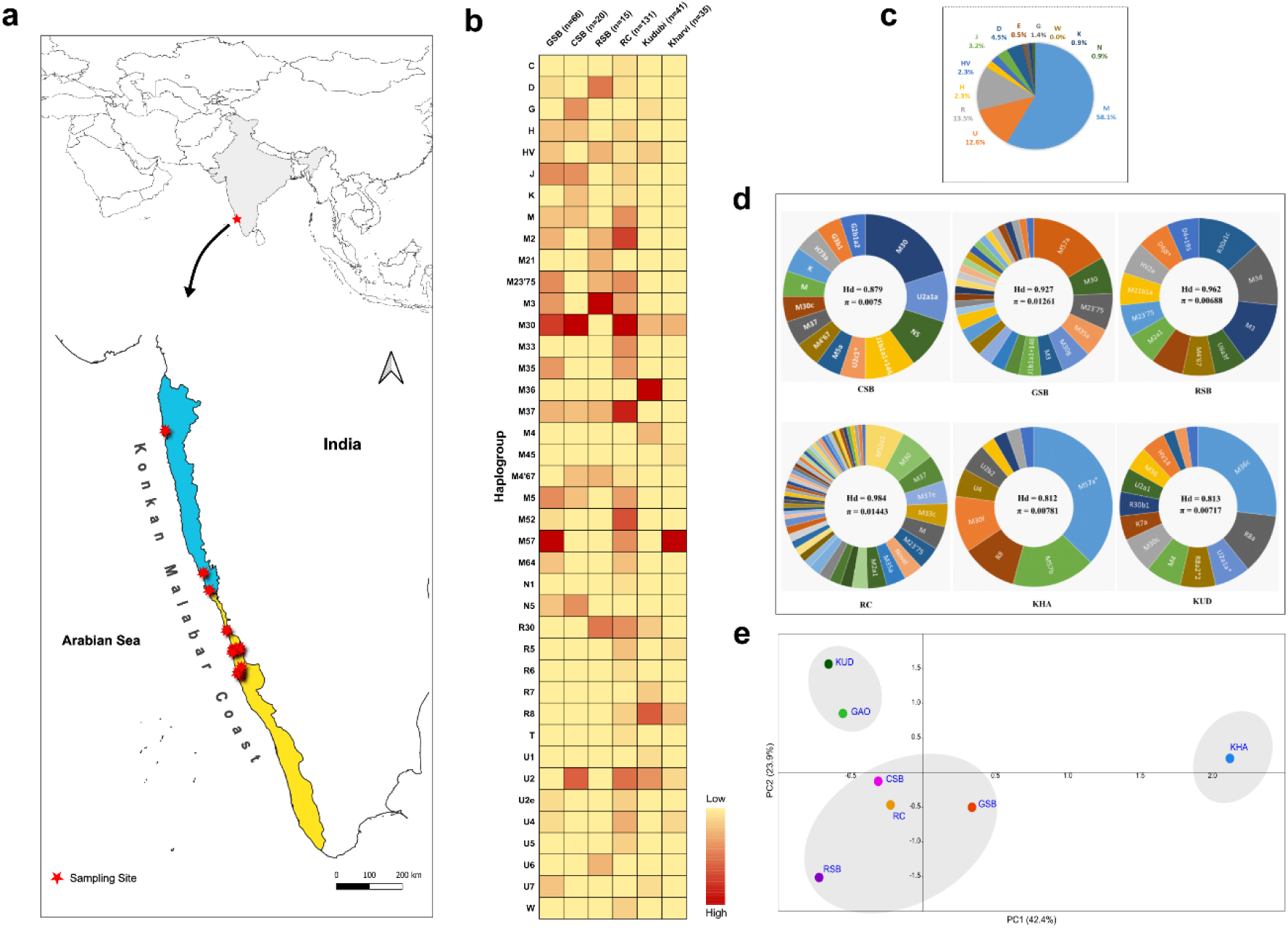
(a) Map showing sampling sites along the Koṅkaṇ-Malabar coast of India. (b) Heatmap of haplogroup frequency distribution in the studied population. (c) Macrohaplogroup distribution in the Koṅkaṇī population. (d) Donut plots for each of the Koṅkaṇī speaking subgroups (e) PCA plot showing clustering pattern in Koṅkaṇī population. Sārasvata groups shown in the plot are Gauḍa Sārasvata Brahmins (GSB), the Citrapur Sārasvata Brahmins (CSB) and Rājāpur Sārasvata Brahmins (RSB) and non-Sārasvata groups are Khārvi (KHA), Kuḍubi (KUD), 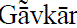 (GAO) and Roman Catholics (RC).

**Fig. 2.** Heatmap showing average pairwise differences between Koṅkaṇī and other populations

In addition to this linguistic dialect geography, Koṅkaṇī speakers distinguish distinct lineages. This main distinction lies between the Sārasvata and non-Sārasvata Koṅkaṇī groups. Sārasvata represents one of five Gauḍa Brahmin groups mentioned in the Sahyādrikhaṇḍa section of the Skandapurāṇa (da Cunha 1881, Deshpande 2010). Koṅkaṇī speakers identifying themselves as being of Sārasvata lineage include the Gauḍa Sārasvata Brahmins (GSB), the Citrapur Sārasvata Brahmins (CSB) and Rājāpur Sārasvata Brahmins (RSB). Sārasvata ancestry is also claimed, however, by some non-Koṅkaṇī speaking Brahmins in western and northwestern India. Endogamous Koṅkaṇī speaking communities who are regarded as being of non-Sārasvata ancestry include Khārvi, Kuḍubi, 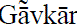, Pāgi, Roman Catholics, Veḷip, Koṅkaṇ Marāṭhā, Daivajña Brahmins. The Koṅkaṇī speaking communities of foreign origin like the Siddis and the Navāyats are also a part of this group.

The origin of Sārasvat lineages is popularly held to have lain on the banks of the Sarasvatī, a mythical river assumed to have existed in the Harappan landscape before its drying. Virendra Nāth Miśra (Miśra 1984; Misra 1994) recapitulates the evidence for the archaeologically well-supported hypothesis that the Sarasvatī river mentioned in the R̥gveda can most likely be identified with the large and now dry riverbed of the Ghaggar-Hākrā, along which hundreds of sites have been discovered. The desertification of the previously flourishing landscape in the interfluve between the Indus and Ganges basins was caused by the weakening of the monsoons and therefore of monsoonal-fed rivers in this region (Giosan et al. 2012). Resultant hydroclimatic stress compelled populations to migrate. Present-day Sārasvats living on the Koṅkaṇ coast believe that their ancestors migrated from the north, and some claim a genealogical affiliation with Kashmiri Paṇḍits.

Against this ethnolinguistically complex background, interdisciplinary research on Koṅkaṇī language and the genomics of its speakers promises to provide a deeper understanding of past migration and present-day settlement. Apart from an unpublished genomic study (Pai 2007) and a recent study on one of the groups speaking Koṅkaṇī language, the Roman Catholic Brahmins (Kumar et al. 2021), most of the work so far is based on historical records, linguistics and anthropology. Using whole mitochondrial DNA of 85 samples and the hypervariable regions (HVS-1 and HVS-2) of 210 samples, we try to understand the maternal ancestry landscape in the Koṅkaṇī population on the south west coast of India.

## Materials and Methods

### Sample collection

Blood samples were collected from 210 volunteers as per the guidelines of Institutional Human Ethics Committee, Mangalore University (MU-IHEC-2020-3 dated 21.08.2020). 2-3 ml blood was collected from unrelated healthy individuals residing along the Koṅkaṇ-Malabar coast of India (Fig. 1a) after obtaining informed written consent. All the donors were personally informed, and their ethnic history was documented. Blood sample were collected and processed as per the guidelines of the Ethics Committee and the Declaration of Helsinki.

Samples were randomly collected at the sample sites (Fig.1) with prior knowledge of the abundance of Koṅkaṇī speakers. In order to reduce sampling errors, detailed discussions were held with the volunteers to know their lineage at least up to 3-4 generations. Sample includes, both Sārasvata (Gauḍa Sārasvata Brahmins (GSB, n=), the Citrapur Sārasvata Brahmins (CSB) and Rājāpur Sārasvata Brahmins (RSB)) and non-Sārasvata groups (Khārvi, Kuḍubi, 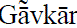, Roman Catholics).

### Genotyping

Genomic DNA was extracted using the standard phenol chloroform method and quantified using Nano Drop spectrophotometer. Mitochondrial DNA (mtDNA) D-loop was amplified for 210 individuals using the primers; 15 F/R, 18 F/R, 22F/R, 23 F/R and 24 F/R primers (Rieder et al. 1998; Mustak et al. 2019) and ABI Veritii (Applied Bio Systems) Thermocycler. PCR conditions used were as follows: initial denaturation at 95 °C for 5 min and 35 cycles of denaturation at 95 °C for 1 min, annealing 52 °C for 30 s, extension at 72 °C for 2 min and a final extension at 72 °C for 5 min. The plates were stored at 4°C until Sanger sequencing. Hypervariable regions (HVS-1 and HVS-2) of mitochondrial DNA (mtDNA) are highly polymorphic, maternally inherited; hence used for population studies (Van Holst Pellekaan et al. 1998; Forster et al. 2002; Maruyama et al. 2010; Guba et al. 2011). Besides these hypervariable regions, for better resolution and divergence time estimation, we used complete mitogenomes from Koṅkaṇī people and compared them with ancient and modern mitogenomes. Whole mitogenomes (n=85) were sequenced using the primers listed in Supplementary Table S1.

## Data analysis

### Estimation of mitochondrial DNA diversity

The sequences were aligned against the revised Cambridge Reference Sequence (rCRS) (Andrews et al. 1999) using the MAFFT algorithm in UniPro Ugene software (Okonechnikov et al. 2012). The variations were listed and run into Haplogrep 3.0 (Schönherr et al. 2023) for haplogroup assignment. Haplogroup frequency distribution from 210 was used to calculate diversity and intra-population structure. DnaSP6 (Rozas et al. 2017) was used to measure nucleotide and haplotype diversity.

### Haplotype sharing and population structure

To understand the extent of haplotype sharing between subgroups, a median joining network (Bandelt et al. 1999) was plotted using the hypervariable region in PopART (Leigh and Bryant 2015). Principal Component Analysis (PCA) plot for intra-population and inter-population analysis was plotted, using PAST 4 software (Hammer et al. 2001) and *prcomp* package in R software (R Core Team 2021) respectively. Variance between the groups and populations within the groups was measured with AMOVA using Arlequin 3.5 software (Excoffier and Lischer 2010). Two datasets were generated by pooling all the available M and N lineage mitogenomes, respectively, from the Indian subcontinent and neighbouring regions to understand the lineage specific affinity in Sārasvata and non-Sārasvata subgroups. Data from previous studies on the Koṅkaṇī population (Kumar et al. 2021), other populations (Maca-Meyer et al. 2003; Metspalu et al. 2004; Baig et al. 2004; Palanichamy et al. 2015, 2004; Sharma et al. 2005, 2018; Thangaraj et al. 2005, 2010; Barik et al. 2008; Chandrasekar et al. 2009; Shah et al. 2011; Barbieri et al. 2012; Pemberton et al. 2012; Pijpe 2013; Ranaweera et al. 2014; Ranasinghe et al. 2015; Larruga et al. 2017; Cabrera et al. 2018; Mustak et al. 2019; Sylvester et al. 2019; Rahman et al. 2020, 2021; Singh et al. 2021) and ancient DNA data (Ehler et al. 2019) were collected and used for this purpose. This dataset included 5128 whole mitogenomes. Nei’s distance matrix was generated using dartR v2 package (Gruber et al. 2018; Mijangos et al. 2022) and visualised in R software.

### Coalescence age estimation and Bayesian Skyline plots

BEAST v2.7.4 was used to do Bayesian age estimation and to generate Bayesian Skyline Plots (BSP) (Bouckaert et al. 2019). Complete mitogenomes were filtered based on the haplogroup they were assigned to. For phylogenetic tree calibration, ancient mitogenomes from AmtDB that had better coverage and C14 dates were employed (Ehler et al. 2019) as tip dates. As an outgroup, the L2c2 mitogenome of a Moreno person (PaMOR16007) was used. HKY substitution model (gamma-distributed rates plus invariant sites) with a strict clock was chosen as per Brandini et al. 2018. For the HVS1 and HVS2 regions, rigorous molecular clocks with mutation rates of 1.292 and 0.369 mutations/site/million years, respectively, were used. Coding region partition was run considering a mutation rate of 0.03 mutations/site/million years (Connell et al. 2022). We have considered pedigree derived mutation rates acknowledging the endogamy that is followed in India even today. We conducted Chain Monte Carlo (MCMC) runs consisting of 100,000,000 steps, with parameters sampled every 10,000 steps. The initial 10% of the steps were discarded as burn-in. Coalescence Bayesian Skyline was selected as the tree prior. BEAUti from the BEAST package (Bouckaert et al. 2019) was used to set the model and parameters. The convergence of MCMC was assessed using the effective sample size (ESS) in Tracer v1.7.2, with ESS > 200 being acceptable. The same software was used to generate Bayesian Skyline Plots (BSP) and a combined BSP was visualised using an in-house R script. For phylogenetic tree construction, a consensus tree was generated using Treeannotaor. The median height coalescent age estimates were plotted using FigTree v1.4.4.

## Results and Discussion

### Maternal ancestry landscape in Koṅkaṇī population

A distinct stratification, observed in our mtDNA data, shows high frequency of M derived haplogroups (58.1%) in the Koṅkaṇī population, followed by other haplogroups, including R (13.5%), U (12.6%), D (4.5%), J (3.2%), HV (2.3%) and H (2.3%) (Fig. 1c). This dominance of M haplogroup in the Indian subcontinent is not surprising, as it was observed in earlier studies (Cordaux et al. 2003; Kivisild et al. 2003; Metspalu et al. 2004).

Haplogroup distribution pattern and genetic affinity for each of the Koṅkaṇī subgroups is as follows,

### Citrapur Sārasvata Brahmins (CSB)

The haplotype diversity (Hd) index for CSB was 0.879 and nucleotide diversity (π) index was 0.0075 (Fig. 1). Subclades of maternal haplogroup M, N, J, G, K, U2 and H are present in CSB. M30 is found abundantly at 25%, followed by U2a1a (10%), N5 (10%), J1b1a1+146 (10%) and G (10%). Haplogroup M30 is Indian-specific, which is ubiquitously found in castes and tribes. The estimated age of this lineage is 15.4+/-6.3 thousand years (Thangaraj et al. 2006). The presence of N5 suggests Mesolithic movement within India, as this lineage is about 35,000 years old in contrast to M and R, which have much older deep lineage in India (Fregel et al. 2015). Haplogroup J is a Middle-Eastern, while U2 is an Indian-specific lineage (Palanichamy et al. 2015). Haplogroup G is found abundantly in East Asian populations (Jin et al. 2009). The presence of H and K haplogroups in CSB suggests European admixture.

### Gauḍa Sārasvata Brahmins (GSB)

The GSB is more diverse group among the Sārasvata Brahmins, with Hd = 0.927 and π = 0.01261 (Fig. 1). Subclades of maternal haplogroup M, N, D4, J1, U2, U7 and HV are present in GSB individuals. Haplogroup M57a is the most abundant subclade, existing at 18.1%. The presence of HV suggests traces of West Asian admixture (Shamoon-Pour et al. 2019). The HV6 subclade is found in other South Indian populations as well (Palanichamy et al. 2015). Haplogroup J originated in the vicinity of the Fertile Crescent and spread across the Europe. The J1b subclade found in GSB is more prevalent in the Romani groups (Malyarchuk et al. 2006; Mendizabal et al. 2011). The U2e1h, U4b1a1a1 and U7 subclades found in this population are West Eurasian (Derenko et al. 2014). Moreover, sub-clusters of U7 are found in high frequencies in India (Palanichamy et al. 2015). Traces of the D4 subclade, common in Central and Northeast Asia, were also found in this population.

### Rājāpur Sārasvata Brahmins (RSB)

In RSB, the haplotype diversity was 0.962 and the nucleotide diversity was 0.00688. MtDNA analysis of RSB population shows the presence of M, R, HV, U and D4 haplogroups (Fig. 1). The most frequent haplogroups found were M3 (13.3%) and M3d (13.3%). Northwest India is highly concentrated with M3 (Metspalu et al. 2004) and therefore the presence of this lineage in RSB hints at its northwestern maternal gene pool. Along with other Indian-specific M subclades, R30a1c (13.03%) was also found in this group. Interestingly, the U6a3f subclade individual was found in this community. U6a is an ancient haplogroup, widespread in North Africa (Secher et al. 2014). The coalescent age of U6a3f (∼6500 years) and its absence in the Indian subcontinent suggest a recent intrusion of this clade into the RSB group, possibly as an outcome of trade or colonization. The presence of West Eurasian haplogroup HV2a, East Eurasian D4 and Indian-specific M3 subclades indicates that the maternal gene pool could be a residue of the Late Pleistocene demographic expansion suggested by Metspalu et al. (2004).

### Roman Catholics (RC)

A recent study on Catholic Brahmins concluded that Roman Catholics come from the same lineage as the Gauḍa Sārasvata Brahmins (Kumar et al. 2021). A continuous religious conversion that took place in the 16^th^ century in Goa has resulted in the formation of this group. It is said that this group along with the GSB community migrated southwards and settled along the Koṅkaṇ Malabar coast. Whilst our results are in line with the findings of Kumar et al. (2021), we find more diversity in Roman Catholics (58 haplogroups) compared to GSB (33 haplogroups), with Hd = 0.984 and π = 0.01443 (Fig. 1). Both Indian and West Eurasian mitochondrial haplogroups are present in Roman Catholics. The most prevalent subclades are M52a1 (7.6%), M30 (6.9%), M37 (5.3%), M37e (4.6%), M33c (4.6%) and M23’75 (4.6%). The basal lineage M (4.6%) is also present in this group. The major difference between GSB and Roman Catholics is the increased number of Indian-specific haplogroups in the latter. The presence of subclades of R30, R5, R6, R8, U2a, U2b and U2c in addition to the West Eurasian (U2e, U4, U7, U5 and HV), African (N1), West Asian (J1, W), Near Eastern (T), Central Asian (C) and East Asian (D) subclades highlights the diversity in this group. This diversity can be attributed to the influence of Christianity, which does not promote caste-based endogamy.

### Khārvi (KHA)

Koṅkaṇī Khārvi is a fishing community found throughout the Koṅkaṇ coast. Presently, the Khārvi are concentrated in coastal towns of Udupi district. Their mitochondrial genome is less diverse (Hd = 0.812 and π = 0.00781), with M57, M57a* and M57b haplogroups accounting to 57% of the total frequency distribution (Fig. 1). These haplogroups are found in GSB and Roman Catholics as well. Haplogroups M30f and R8 were found at 11.4%, whereas U2b2, U4, R8, R5 and M45a were present in small trace amounts. Low haplotype diversity and nucleotide diversity indicate a recent population bottleneck or a founder event in the maternal lineage of this group (Grant and Bowen 1998). Probably this group was a part of the tribes that settled in the Western Indian belt, moved southward and adopted marine fishing as their primary occupation. This assumption is based on the presence of M57 in Ḍoṅgrī Bhil, Kātkarī (Kathauḍī) and Kaṭhākur tribes from western India. The TMRCA age for this haplogroup is estimated to have lain around 45,000 and 25,000 years ago (Chandrasekar et al. 2009). The presence of the R8 haplogroup without any subclades at a relatively moderate frequency again hints at a recent founder event.

### Kuḍubi (KUD)

The diversity indices for Kuḍubi population are Hd = 0.813 and π = 0.00717 (Fig. 1). The subclades M36c (26.8%) and R8a (12.2%) mainly contribute to the mitochondrial haplogroup diversity in this group. The abundance of only two sub-lineages suggests a founder event in this population. In addition, Indian-specific subclades (M, R, U2), West Eurasian subclades (U1, U7, HV) and East Asian subclades (G2a5) are present in this population. Haplogroups M36 and M36c have been found earlier in Dravidian groups of South India (Chandrasekar et al. 2009). The subclades R8a, R8a2*2 and R7a are frequent amongst Austroasiatic groups (Thangaraj et al. 2009). The R8 subclades are also present in Mālekuḍiyā tribe in Karnāṭaka (Sylvester et al. 2019). Based on the TMRCA of M36 (∼34,000 to ∼26,000 years) and R8a (∼40,600 years), we can conclude that this group has admixed with both the Dravidian and Austroasiatic gene pools. The incidence of East Asian G2a5 could be attributed to a recent admixture. HV14 originated in Iran and migrated to South Asia before and during the Neolithic period (Shamoon-Pour et al. 2019). While U1 is found in Eastern Europe, Anatolia and the Near East, U7 is West-Eurasian-specific (Metspalu et al. 2004). U1 haplogroup was recently reported to be a tracer dye for the maternal link between the Koraga tribe and the north western region (Sequeira et al. 2024b). The Kuḍubi tribe has a strong presence along the west coast and may have intermixed with neighboring tribes.

We performed PCA based on the haplogroup frequencies of subgroups within the Koṅkaṇī speaking population and found three distinct clusters (Fig. 1e). One of the clusters included the Kuḍubi and 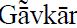 groups. Traditionally these groups are dependent on agriculture. The second cluster consists of Sārasvata group and Roman Catholics. Except for Roman Catholics, who are considered offshoots of GSBs (Kumar et al. 2021), and are now a heterogeneous group, this cluster includes non-agrarian communities that occupy positions of authority, trade and land property. Khārvi, a traditionally fishing group clustered away from the other Koṅkaṇī subgroups. We observed similar population structure in the pairwise F_st_ matrix (Supplementary Fig. S1).

To understand the lineage sharing patern within the Koṅkaṇī population, a median joining network of haplotypes was constructed (Supplementary Fig. S2). Three major clusters of haplotypes responsible for the observed variation in GSB, Khārvi and Kuḍubi were found to form a star-like network, implying that these haplotypes arose from separate founders. The interlinking between the clusters suggests maternal lineage sharing between these groups. More than 50% of these variations in the subgroups of Koṅkaṇī population is retained, probably due to genetic isolation. Interestingly, the cluster with Kuḍubi and GSB haplotypes does not contain haplotypes present in Khārvi. This observation when considered along with the lowest F_st_ values for Khārvi, Ḍoṅgrī Bhil and Andamanese suggests that the maternal gene pool of this group formed at a different time depth. We observe intermixing between GSB and Khārvi that could be attributed to their occupational exchange. Both these populations are involved in trade and, therefore, concentrated at port towns.

### Affinities between Koṅkaṇī and other populations

To understand their overall variation, AMOVA using haplotype frequencies was performed against different populations based on their geographical distribution. The overall amount of variation between geographically identified population groups was low. The highest percentage variation was observed between Sārasvata and Andamanese groups (19.29%), which was much higher when compared to the non-Sārasvata groups (11.53%). This finding suggests that while the maternal gene pool of both Sārasvata and non-Sārasvata groups was similar to the maternal gene pool of the Indian subcontinent, the former underwent a greater number of admixture events before attaining its present genetic make-up. Another major difference was the higher variation observed between the non-Sārasvata (2.54%) and the Western and Central Indian populations when compared to the Sārasvata (0.42%). This finding is in line with the F_st_ results, which show affinity between the maternal roots of these groups. A similar interpretation can be made for Northern and Northwest Indian groups as well. The AMOVA results allow us to conclude that the negligible variation found between the subgroups of the Koṅkaṇī population and other geographically separated Indian populations (except the Andamanese) was mostly due to the basal clades that remained in the founder populations that formed in the Late Pleistocene (based on the TMRCA of abundant haplogroups) and subsequently remained isolated.

In order to understand the genetic affinity of Koṅkaṇī population with other populations, we compared the haplogroup frequency distribution of the subgroups with other populations on a PCA plot (Fig. 3). Most of the subgroups clustered closer to a convergence point possibly representing populations with higher admixture components because more unique southern Indian tribal groups clustered at one extreme, while the other extreme consisted of Western Indian and Northwest Indian populations. Both Sārasvata and non-Sārasvata groups clustered separately. A distinct pattern was observed for each of the subgroups of Koṅkaṇī populations in different cluster analysis methods, suggesting that the ancestral maternal gene pool had both unique and admixed components.

**Fig. 3.**
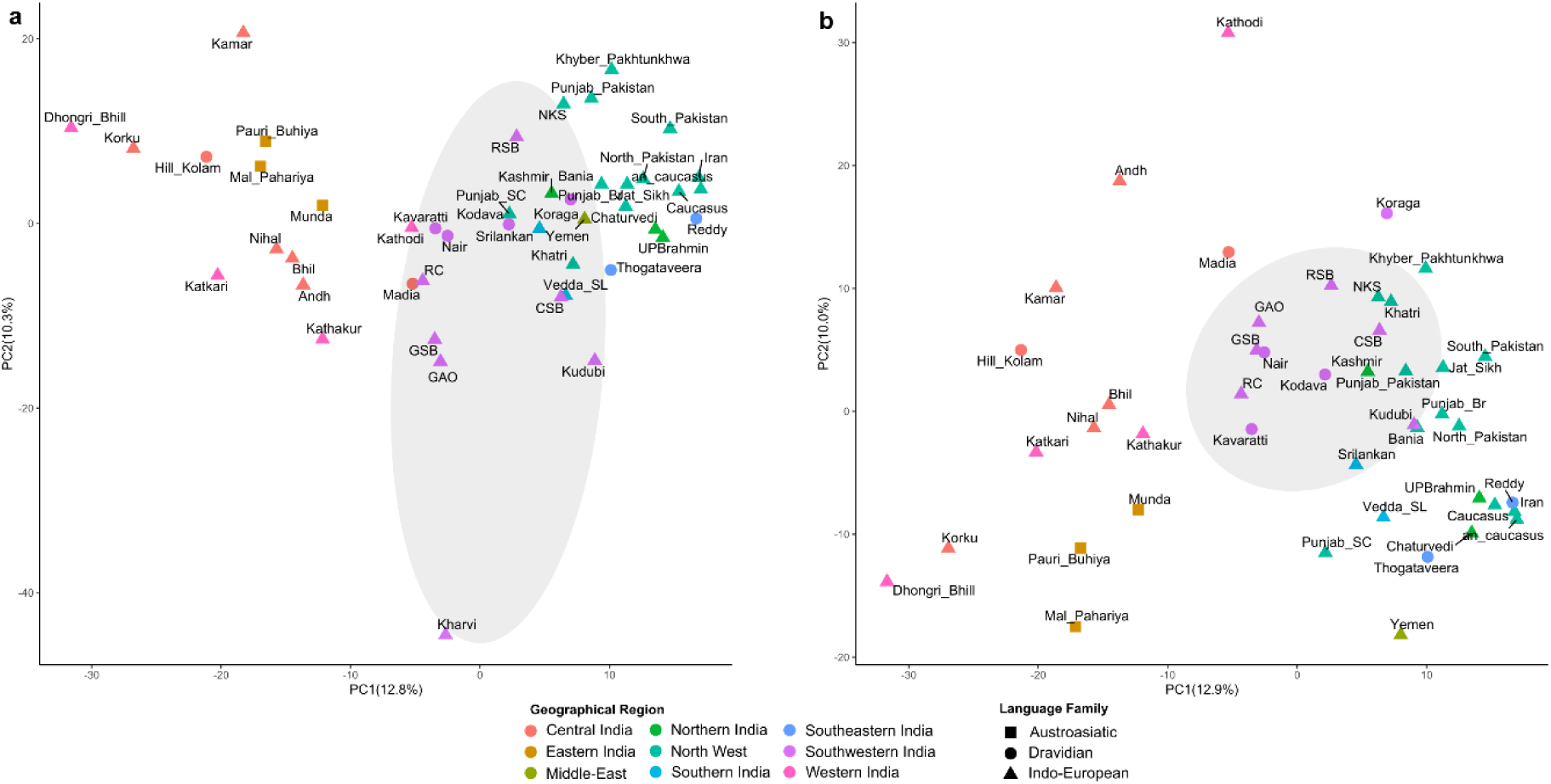
PCA showing clustering pattern between Koṅkaṇī and other populations. (a) with outliers (b) without outliers. Koṅkaṇī subgroups are within the grey ellipse.

### Demographic reconstruction of major lineages in Koṅkaṇī population

In order to understand the effect of external factors like mass migration, climatic changes and geographical barriers on the maternal genetic landscape of Koṅkaṇī population, we constructed Bayesian skyline plots for major haplogroups (Fig. 4). Irrespective of their social affiliation, we observe a rapid population expansion between 50 and 40 Kya. Hypotheses like “Eurasian population hub” and “delayed expansion” in the Middle East-Near East region have been based on similar observations (Xing et al. 2010; Vallini et al. 2022). An expansion around 50 Kya is also reported in the paternal lineage of eight global regions as well (Karmin et al. 2015).

**Fig. 4.**
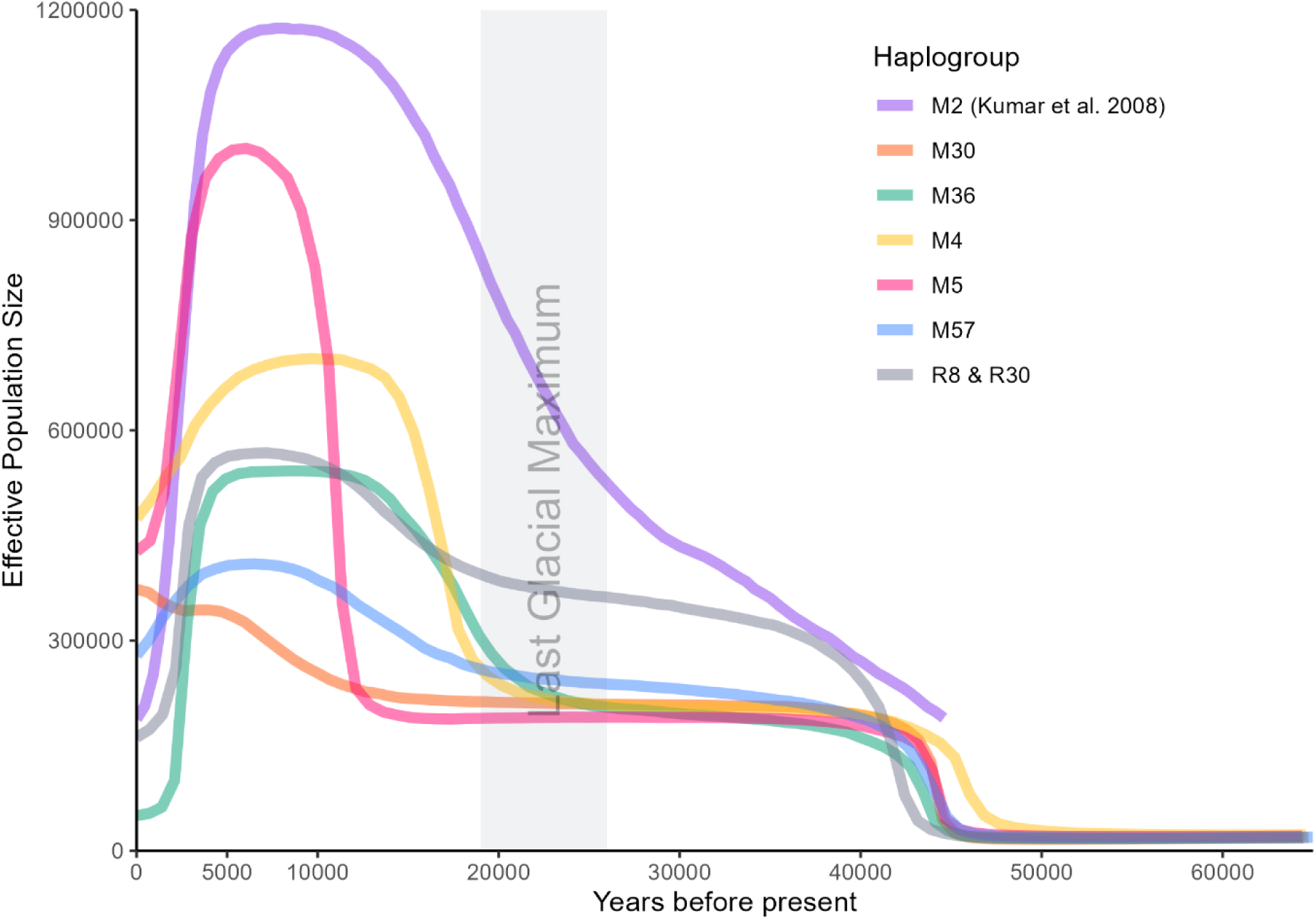
Bayesian skyline plots for major haplogroups in Koṅkaṇī population

The next increase in N_e_ between 20 and 10 Kya is more gradual. Such expansion in the maternal lineages in the time-depth that coincides with the LGM recovery phase, is not universally seen in all autochthonous haplogroups in the Indian subcontinent. In Silva et al., 2017 (Silva et al. 2017), Bayesian skyline plots of M haplogroup varied distinctly between West, Central, South and East regions. Similarly, in Kumar et al. (2008) (Kumar et al. 2008), BSPs of M2 haplogroup showed variation among different regions. We performed Bayesian analysis of M2 mitogenome data generated in Kumar et al. (2008) (Kumar et al. 2008) using the parameters described in the Materials and Methods section. The gradual rise in the effective population size that began before the LGM continued until the Holocene where we see a drastic dip in the last 5000 years. M2 lineage flourished since the earliest migrations out of Africa and was largely unaffected by the LGM.

The dip in N_e_ observed in most of the haplogroups in the last 5000 years can be attributed to the stringent social stratification. Interestingly we have not observed such a dip in M30. This may be due to the ubiquitous presence of this haplogroup in castes and tribes (Supplementary Fig. S10). Overall, the Bayesian analysis indicates that both the Last Glacial Maximum (LGM) and the recent caste-based stratification have had a greater impact on the maternal ancestry of the Koṅkaṇī population than the expansion of Neolithic agriculture.

Phylogenetic trees were constructed for major haplogroups using Bayesian method implemented in the Beast v2.7.4. All annotated trees can be found as Supplementary Fig. S4-S11. Apart from U1 clade found in Kuḍubi and U8 found in the CSB, which show an estimated TMRCA of 34-38000 and 32000 respectively, all other clusters are dated around the LGM and LGM recovery phase. Our estimates are similar to the divergence times for Indian clusters reported in earlier studies (Supplementary Table S3).

### Influence of macrohaplogroup N lineages on the maternal ancestry of west coast populations

The average pairwise F_st_ values permit us to interpret the divergence of two populations from each other having disturbed the Hardy-Weinberg equilibrium. We first performed haplogroup frequency based Fst analysis using Konkani and published datasets. The Sārasvata subgroups displayed a similar pattern of divergence compared to the non-Sārasvata subgroups (Supplementary Fig. S3). The Khārvi community showed the lowest F_st_ values with the Önge (0.851) and Greater Andamanese (0.777) populations, followed by the west Indian populations such as the Ḍoṅgrī Bhil (0.965) and Nihal (0.985). Although these results indicate deep lineage relationship between the mainland populations and the Andamanese tribes, we performed sequence variation based Fst analysis in anticipation of similar results. Nei’s distance matrix was generated using 418 D-loop sequences which included Koṅkaṇī samples (Supplementary Fig. S12). Kuḍubi and Roman Catholics clustered with the ancient Iberian and Mediterranean samples, whereas Khārvi, which showed lesser pairwise differences with the Andamanese based on haplogroup frequency distribution, clustered with the Sārasvata group. These results highlight the role of lineage-specific variants in diversifying the maternal ancestry of Koṅkaṇī population.

Hence, we performed a similar analysis using all the available complete mitogenomes belonging to macrohaplogroup M and N lineages separately. Nei’s distance matrix for M lineage was based on 800 mitogenomes and N lineage consisted of 4328 mitogenomes. Both the datasets included ancient mitochondrial sequences. As expected, we observed distinct clusters in both the lineages (Fig 5). In the N lineage, we observed a more scattered distribution of the Koṅkaṇī subgroups compared to their distribution in the M lineage. Kharvi and Kudubi cluster together with the other south-western and north-western populations. Interestingly, GSB also belonged to this cluster sharing the clade with Kalash population of Pakistan. CSB and RSB clustered with the central Indian Sahāriyā (Cakāriyā) tribe and ancient Altai population. An earlier study reports the oldest paternal R1a lineage in the Sahāriyā tribe (Sharma 2009). In addition, this nomadic tribe and the tribes of the Altai region show similar levels of R1a haplogroup. Now the question is whether the maternal N lineages found in this cluster have any links with the paternal R1a haplogroup? A comprehensive analysis performed by us using Y-chromosome STR markers shows a similar clustering pattern between the Sārasvata Brahmins and the Altaians in the R1a lineage (Sequeira et al. 2024a). These findings indicate that this admixture was not male mediated, and may have resulted in the assimilation of West-Eurasian N lineages in the Sārasvata group.

**Fig. 5.**
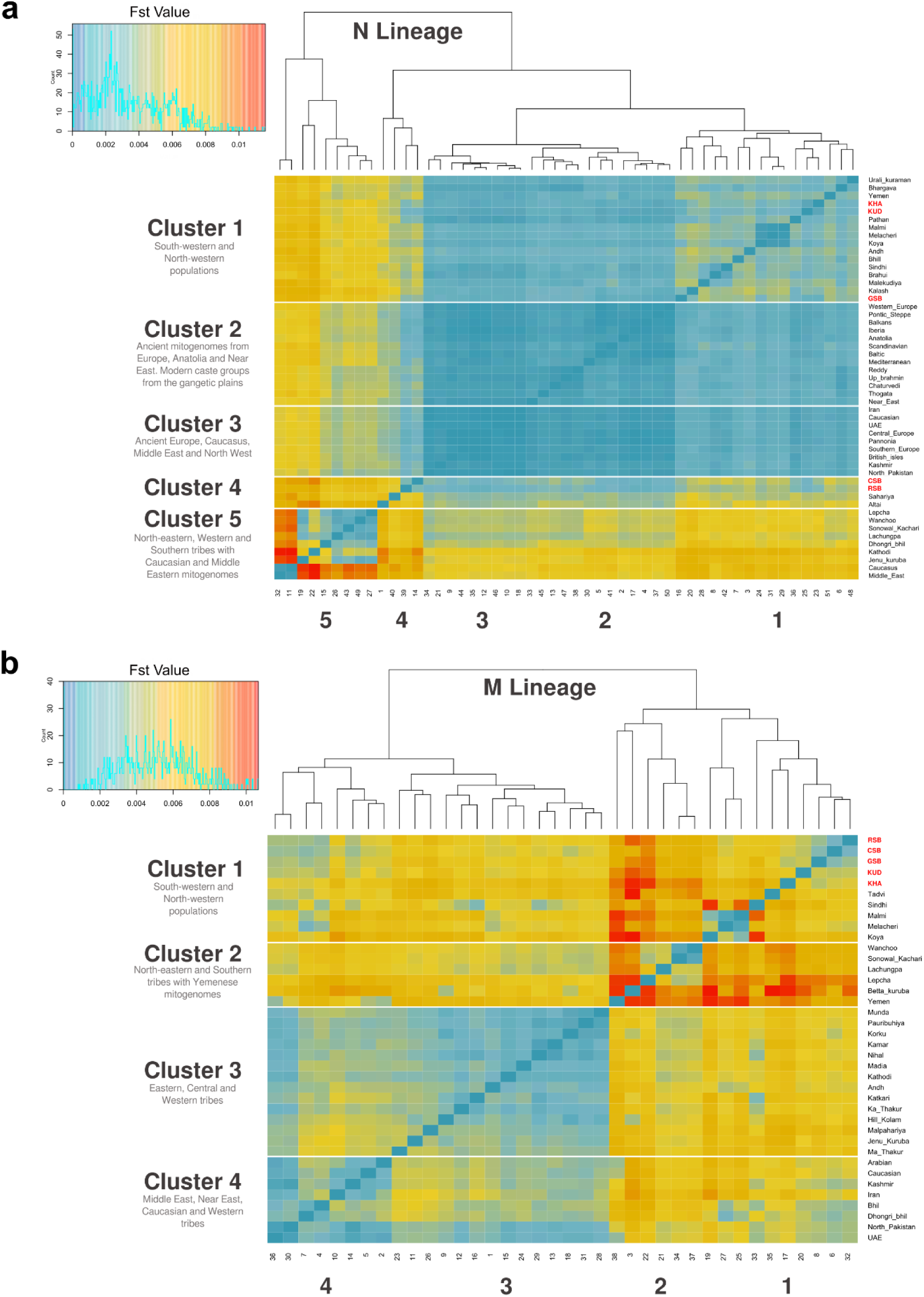
Heatmap showing Nei’s distance matrix for N lineage (A) and M lineage (B). Koṅkaṇī subgroups are shown in red font. Clusters correspond to the major clades observed in the phylogram.

In the M lineage cladogram, the Sārasvata group clustered together. Although Kuḍubi belonged to the same cluster, it shows greater genetic distance from other Sārasvata members. Interestingly, Khārvi population clustered with another western tribe called Taḍvī. Overall, most of the Koṅkaṇī subgroups showed greater affinity with the Western Indian tribes like Bhil and Korku, and UAE, North Pakistan and Kashmir populations. Kharvi population was an exception to this trend. Two broad clusters within the M lineage, one more diverse and the other more similar, suggests that the former cluster has undergone higher levels of admixture compared to the latter which includes most of the ancient Indian tribes. It is interesting to note that the measure of genetic distance between them is such that the Koṅkaṇī population shares the clade with the north-eastern populations.

What stands out is that the diversity of Koṅkaṇī subgroups is dictated more by the N lineages compared to the M lineages found in them. From the estimated TMRCA (Supplementary Table S3), it is evident that these N lineages are ancient going back to the Last Glacial Maximum era. In order to understand the effect of the LGM on both M and N lineages, we performed PCA using the haplogroup frequencies of populations of the Indian subcontinent. Published whole mitogenome data of 3054 samples from 58 populations including the Koṅkaṇī were analysed using Haplogrep software and haplogroups were reassigned. The pre and post LGM haplogroup frequency variation in these populations were visualised on PCA plots. The scatter was distinctly different in both the plots (Fig. 6 a and b). In order to measure the contribution of haplogroups responsible for the scatter pattern, a biplot was constructed using the same principal components (Supplementary Fig. S13). In the pre-LGM era, haplogroup M2 and R30 formed two distinct clines (Fig. 6 a). The third was composed of haplogroups like N5, M31 and M32 found in Sahāriyā and Andaman tribes. The M2 cline consisted of Bëṭṭʉ Kuṟumba and most of the Western Indian tribes. Whereas the R30 cline included Ūrāḷi Kuruman and the Lakshadweep population. An earlier study assigned R8 haplogroup to all the R30 samples (reassigned here) from the Ūrāḷi Kuruman tribe (Sylvester et al. 2019). This ambiguity has been highlighted in a recent study on Kavaratti Island of the Lakshadweep (Lakṣadvīpŭ) or Laccadive archipelago (Tayyeh et al. 2023). Nevertheless, the scatter does show three extremities on a more compact pattern compared to that of the post-LGM era plot (Fig. 6 b).

**Fig. 6.**
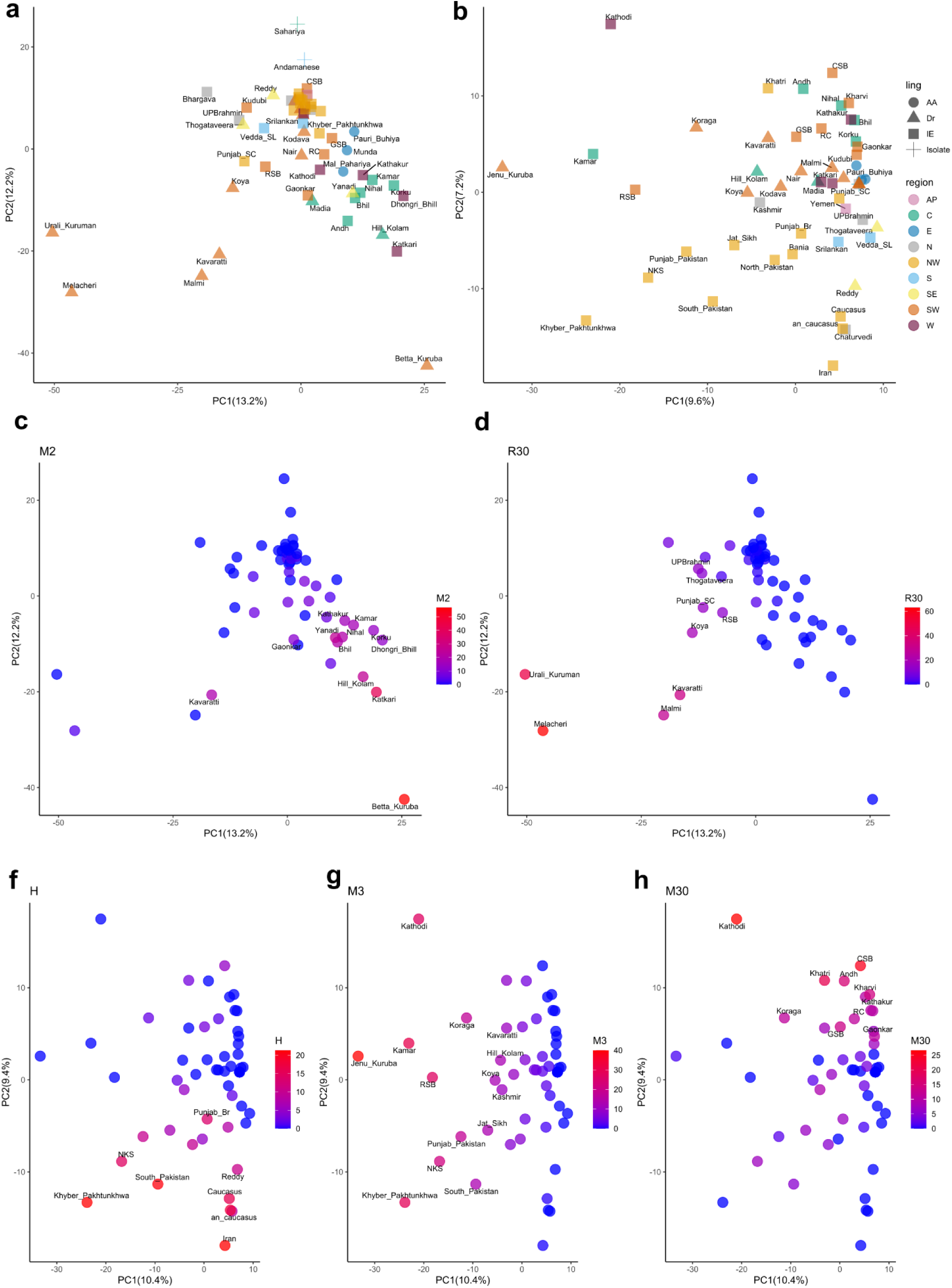
Principal component analysis using frequencies of haplogroups from the pre LGM (a) and post LGM era (b). The contribution of specific haplogroups to the overall variance was visualised using a biplot (S1 Fig. 16). Frequency distribution of these putative founder haplogroups in the pre LGM (c and d) and post LGM (e, f and g) era is shown in the same PCA plot.

In the post-LGM PCA plot, at least three haplogroups dictate most of the variance. The Caucasian/Near Eastern cluster shows higher frequency of H haplogroup. Jēnu Kuṟumba and most of the western Indian populations show higher frequencies of M3 haplogroup. A third cluster which includes Koṅkaṇī and other western populations is predominated by M30 haplogroup. Through this analysis we find that the pre-LGM gene pool of the Indian subcontinent although diverse, was more homogenous. The maternal gene pool shows that the founder effect observed in some western and southern Indian tribes dates back to the pre-LGM era. Paternal haplogroup F and H, which are found at high frequencies in these tribes originate around the same time depth, (ArunKumar et al. 2012) suggesting that these early migrations are a part of human movement through the north-western region subsequent to the earliest Out-of-Africa dispersal. In the post LGM era, the heterogeneity may be attributed to multiple stints of migrations throughout the Holocene. Interestingly, most of it is observed on the western part of the Indian subcontinent. Although it may seem obvious that agricultural expansion and late bronze age migration caused such heterogeneity in the maternal gene pool of the western front, it is very unlikely that primitive hunter gatherers like Jēnu Kuṟumba, Koraga and Kātkarī (Kathauḍī) shared the same story. A more plausible explanation for post-LGM but pre-Holocene heterogeneity in the maternal gene pool of the western coast is influx of tribes from the north-western/near eastern region in the LGM recovery phase when most of the coastline was exposed due to lower sea levels (Lambeck et al. 2014).

In the case of Koṅkaṇī population, the movement of Sārasvata females from the north-western region as well as intake of females from local populations may have resulted in a more homogenous M lineage irrespective of their social affiliation. However, in the N lineage, although we see a north-western continuity in both Sārasvata and non-Sārasvata gene pool, the time depth of such continuity varies. The clustering of Kuḍubi with ancient Iberian samples is evidence for the post LGM continuity whereas GSB clustering with Kalash is relatively recent (Ayub et al. 2015). The cluster including CSB, RSB, Sahāriyā and ancient Altai may represent a residue of the Steppe maternal ancestry.

## Conclusion

The genetic landscape of the Koṅkaṇī population is an important piece of the puzzle in the documentation of the ethnolinguistic history of the western coast of India. Our findings on the maternal ancestry of this population help to augment our understanding of the prehistoric demographic dynamics of this region. Although the time depth of maternal components tends not to fall primarily in the more recent epoch of civilisation and reconstructible language family relationships, the maternal landscape certainly provides insights about the aboriginals of this region and their adaptations. Here, we report the mitochondrial genomic diversity within the Koṅkaṇī population. Based on our analyses, we suggest that there was active movement in this region in the Late Pleistocene and that the bottleneck events that happened then and through the Holocene are responsible for the clustering patterns that can now be discerned in the maternal gene pool of the Koṅkaṇī population. The Last Glacial Maximum has played an important role in shaping the maternal ancestry of many indigenous populations in the Indian subcontinent (Silva et al. 2017). The same phenomenon is one of the events responsible for present day landscape of maternal lineages in Europe (Richards et al. 2000; Jones et al. 2015; Posth et al. 2016; Fernández-López de Pablo et al. 2019; Bortolini et al. 2021). In the maternal lineage of the Koṅkaṇi population, we have discovered both pre– and post-Last Glacial Maximum (LGM) haplogroups. Upon studying individual subpopulations, we have identified a distinctive distribution pattern in both Sārasvata and non-Sārasvata groups. A major proportion of the maternal diversity comes from ancient N lineages that originated from a diverse ancient source population, likely in the Near East region close to the Last Glacial Maximum, and assimilated in these groups due to the Holocene movements.

## Abbreviations

GSB: Gauḍa Sārasvata Brahmins
CSB: Citrapur Sārasvata Brahmins
RSB: Rājāpur Sārasvata Brahmins
KHA: Khārvi
KUD: Kuḍubi
GAO: 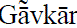
RC: Roman Catholic

## Author Contributions

JJS, MSM and KT conceived the study. JJS collected the samples, performed the experiments and statistical analyses, and wrote the first draft. LK performed additional analysis and improved the manuscript. GvD reviewed, revised and redacted the entire manuscript draft. KT provided research materials. KT and MSM verified the experimental design and contributed to the final draft. All authors approved the submitted manuscript.

## Acknowledgments

JJS is grateful to all the volunteers and staff who participated in this study. JJS and MSM acknowledge the Koṅkaṇī Adhyāyana Pīṭha of Mangalore University and the late Basti Vaman Shenoy, founder of the Viśva Koṅkaṇī Kendra in Mangalore, for their support and guidance. KT was supported by J C Bose Fellowship from the Science and Engineering Research Board (SERB), Department of Science and Technology, Government of India (JCB/2019/000027).

## Data and Code Availability

The datasets generated and/or analyzed during the current study are available from the corresponding author on reasonable request. The sequenced data have been submitted to GenBank under the study accession number PQ306076 – PQ306160.

## Ethics approval

This study was performed in line with the principles of the Declaration of Helsinki. Approval was granted by the Institutional Human Ethics Committee, Mangalore University (MU-IHEC-2020-3 dated 21.08.2020).

## Consent to participate

Written informed consent was obtained from all individual participants included in the study.

## Supplemental Data

Supplementary file1.pdf

Supplementary file2.xlsx

## Web Resources

QGIS, https://qgis.org/en/site/

