## Supplementary_File 1 for "Maternal genetic affinities of Koṅkaṇī population in the southwest coast of India"

**Legend:**

Table S1: Primers used for targeted and whole mitogenome sequencing

Table S2. AMOVA results between population groups based on geographical distribution

Table S3: TMRCA (in years before present) for major haplogroups found in Koṅkaṇī populations

Fig. S1: mtDNA Pairwise Fst matrix for Koṅkaṇī population

Fig. S2: Haplotype network for Koṅkaṇī subpopulations

Fig. S3: Heatmap showing average pairwise differences between Koṅkaṇī and other populations

Fig. S4: M57 phylogenetic tree

Fig. S5: M36 phylogenetic tree

Fig.S6: R8 and R30 phylogenetic tree

Fig.S7: U and K1 phylogenetic tree

Fig.S8: M4 Phylogenetic tree

Fig.S9: M5 Phylogenetic tree

Fig.S10: M30 Phylogenetic tree

Fig.S11: M3 Phylogenetic tree

Fig. S12 Heatmap of Nei's distance matrix based on the D-loop comparing Konkani subgroups with selected populations that showed similarities in haplogroup frequencies.

Fig. S13 Biplots showing the haplogroup frequency contribution to the scatter in Pre and Post LGM era

**About Koṅkaṇī dialects**

The northern Koṅkaṇī dialects comprise the North Goan or Bārdesī variety, upon which standard Koṅkaṇī was primarily based in the latter part of the Portuguese colonial period from the 18th century, and the Central Goan or Antrujī variety of the Phoṇḍā region, upon which Standard Koṅkaṇī has increasingly been based since Goa's accession to the Indian Union. Other northern Koṅkaṇī dialects are Mālvaṇī and Kuḍāḷī, but also include, to the north, the Rājāpurī dialect that since the 17th century has been spoken in Rājāpur in what today is Mahārāṣṭra as well as, to the south, South Canara Christian Koṅkaṇī, which essentially represents a transplanted Bārdesī variety spoken in South Canara since the 18th century.

The southern Koṅkaṇī dialects comprise the South Goan or Sāṣṭī variety, upon which Standard Koṅkaṇī was primarily based during the early part of the Portuguese colonial period from the 15th to the 17th century. Other southern dialects resulting from the southward coastal spread of the Sāṣṭī speech variety during the colonial period include Kārvārī, North Canara Koṅkaṇī and South Canara Hindu Koṅkaṇī, the latter dialect also being spoken in Kāsaragōḍu and Kāññaṅgāḍu in Kerala. The Koṅkaṇī dialect spoken in Cochin (*Kocci*) since the early Portuguese colonial period can be classified as a South Goan or Sāṣṭī variety, though perhaps with some Bārdesī influence.

Navāyatī Koṅkaṇī and Kũḍbī Koṅkaṇī are examples of community-specific Koṅkaṇī sociolects. Navāyatī Koṅkaṇī is spoken by Muslims settled at and around the coastal town of Bhaṭkaḷa in North Canara, whereas Kũḍbī Koṅkaṇī is a label applied to the dialect of Koṅkaṇī spoken along the Kerala coast since the early Portuguese colonial period. Variation can even be heard in the name of the language itself. For example, north of Carwar, the names Koṅkaṇ and Koṅkaṇī tend to be pronounced *Kokaṇ* and *Kokaṇī*, i.e., without the final velar nasal in the first syllable, whereas in Mangalorean Koṅkaṇī the name of the language is pronounced *Koṅkṇī*, i.e., with syncopation of the second vowel, and speakers of the language are referred to as *Koṅkaṇ*, plural *Koṅkṇe*.

**Table S1: Primers used for targeted and whole mitogenome sequencing**

| Primer | Seq 5'-3' | Length | 3' Position | Product (bp) |
|---|---|---|---|---|
| 1F | CTCCTCAAAGCAATACACTG | 20 | 611 | |
| 1R | TGCTAAATCCACCTTCGACC | 20 | 1411 | 840 |
| 2F | CGATCAACCTCACCACCTCT | 20 | 1245 | |
| 2R | TGGACAACCAGCTATCACCA | 20 | 2007 | 802 |
| 3F | GGACTAACCCCTATACCTTCTGC | 23 | 1854 | |
| 3R | GGCAGGTCAATTTCACTGGT | 20 | 2669 | 860 |
| 4F | AAATCTTACCCCGCCTGTTT | 20 | 2499 | |
| 4R | AGGAATGCCATTGCGATTAG | 20 | 3348 | 887 |
| 5F | TACTTCACAAAGCGCCTTCC | 20 | 3169 | |
| 5R | ATGAAGAATAGGGCGAAGGG | 20 | 3981 | 832 |
| 6F | TGGCTCCTTTAACCTCTCCA | 20 | 3796 | |
| 6R | AAGGATTATGGATGCGGTTG | 20 | 4654 | 898 |
| 7F | ACTAATTAATCCCCTGGCCC | 20 | 4485 | |
| 7R | AATGGGGTGGGTTTTGTATG | 20 | 5420 | 975 |
| 8F | CTAACCGGCTTTTTGCCC | 18 | 5255 | |
| 8R | ACCTAGAAGGTTGCCTGGCT | 20 | 6031 | 814 |
| 9F | GAGGCCTAACCCCTGTCTTT | 20 | 5855 | |
| 9R | ATTCCGAAGCCTGGTAGGAT | 20 | 6642 | 827 |
| 10F | CTCTTCGTCTGATCCGTCCT | 20 | 6468 | |
| 10R | AGCGAAGGCTTCTCAAATCA | 20 | 7315 | 886 |
| 11F | ACGCCAAAATCCATTTCACT | 20 | 7148 | |
| 11R | CGGGAATTGCATCTGTTTTT | 20 | 8095 | 987 |
| 12F | ACGAGTACACCGACTACGGC | 20 | 7937 | |
| 12R | TGGGTGGTTGGTGTAAATGA | 20 | 8797 | 900 |
| 13F | TTTCCCCCTCTATTGATCCC | 20 | 8621 | |
| 13R | GTGGCCTTGGTATGTGCTTT | 20 | 9397 | 816 |
| 14F | CCCACCAATCACATGCCTAT | 20 | 9230 | |
| 14R | TGTAGCCGTTGAGTTGTGGT | 20 | 10130 | 940 |
| 15F | TCTCCATCTATTGATGAGGGTCT | 23 | 9989 | |
| 15R | AATTAGGCTGTGGGTGGTTG | 20 | 10837 | 891 |
| 16F | GCCATACTAGTCTTTGCCGC | 20 | 10672 | |
| 16R | TTGAGAATGAGTGTGAGGCG | 20 | 11472 | 840 |
| 17F | TCACTCTCACTGCCCAAGAA | 20 | 11314 | |
| 17R | GGAGAATGGGGGATAGGTGT | 20 | 12076 | 802 |
| 18F | TATCACTCTCCTACTTACAG | 20 | 11948 | |
| 18R | AGAAGGATATAATTCCTACG | 20 | 12772 | 866 |
| 19F | AAACAACCCAGCTCTCCCTAA | 21 | 12571 | |
| 19R | TCGATGATGTGGTCTTTGGA | 20 | 13507 | 977 |
| 20F | ACATCTGTACCCACGCCTTC | 20 | 13338 | |
| 20R | AGAGGGGTCAGGGTTGATTC | 20 | 14268 | 970 |
| 21F | GCATAATTAAACTTTACTTC | 20 | 14000 | |
| 21R | AGAATATTGAGGCGCCATTG | 20 | 14998 | 938 |
| 22F | TGAAACTTCGGCTCACTCCT | 20 | 14856 | |
| 22R | AGCTTTGGGTGCTAATGGTG | 20 | 15978 | 1162 |
| 23F | TCATTGGACAAGTAGCATCC | 20 | 15811 | |
| 23R | GAGTGGTTAATAGGGTGATAG | 21 | 16215 | 765 |
| 24F | CACCATCCTCCGTGAAATCA | 20 | 16420 | |
| 24R | AGGCTAAGCGTTTTGAGCTG | 20 | 775 | 954 |

#### Table S2. AMOVA results between population groups based on geographical distribution

| Group | Source of Variation | % Variation: West Eurasian | African | Northern & North West Indian | Western & Central Indian | Southern Indian | Gangetic | Sinhalese & Maldivian | Andaman |
|---|---|---|---|---|---|---|---|---|---|
| Sārasvata | Among groups | 2.82 | 2.68 | 1.55 | 0.42 | 0.12 | 2.17 | 1.15 | 19.29 |
| | Among populations within groups | 2.73 | 1.33 | 3.01 | 6.57 | 20.56 | 1.85 | 1.69 | 0.52 |
| | Within groups | 94.45 | 96 | 95.44 | 93.01 | 79.32 | 95.98 | 97.17 | 80.18 |
| | FSC | 0.02806 | 0.01362 | 0.03056 | 0.06598 | 0.20582 | 0.01888 | 0.01705 | 0.00649 |
| | FST | 0.05548 | 0.04002 | 0.04559 | 0.06988 | 0.20679 | 0.04018 | 0.02832 | 0.19818 |
| | FCT | 0.02822 | 0.02677 | 0.0155 | 0.00417 | 0.00123 | 0.02171 | 0.01147 | 0.19295 |
| non-Sārasvata | Among groups | 1.25 | -0.48 | 2.7 | 2.54 | -0.19 | 0.94 | -8.88 | 11.53 |
| | Among populations within groups | 5.93 | 8.8 | 5.31 | 7.93 | 26.18 | 9.27 | 13.73 | 11.56 |
| | Within groups | 92.83 | 91.68 | 91.99 | 89.53 | 74.01 | 89.79 | 95.15 | 76.91 |
| | FSC | 0.06001 | 0.08754 | 0.0546 | 0.08134 | 0.26133 | 0.09356 | 0.1261 | 0.13066 |
| | FST | 0.07173 | 0.08315 | 0.08015 | 0.10469 | 0.25992 | 0.10207 | 0.04847 | 0.23092 |
| | FCT | 0.01246 | -0.00481 | 0.02702 | 0.02542 | -0.00191 | 0.00939 | -0.08883 | 0.11533 |

#### Table S3: TMRCA (in years before present) for major haplogroups found in Koṅkaṇī populations

| Haplogroup | Present Study* | [1][#] | [2] | Others |
|---|---|---|---|---|
| M57 | 43,852 | 45,000 | 30,220 | |
| | 19,620 (all Koṅkaṇī) | | | |
| M57a | 16,320 (Khārvi) | 39,000 | 22,506 | |
| M57b | 12,660 (Khārvi) | 3,000 | 14,920 | |
| M36 | 21,000 (Kuḍubi) | 34,000 | 32,094 | |
| M30 | 15,407 (all Koṅkaṇī) | 15,000 | 17,431 | |
| M3 | 16,410 (CSB)<br>13,134 (RSB) | 23,000 | 23904 | 17,300[d] (India) |
| M5 | 9,301 (CSB) | 39,000 | 37,067 | 28,200[b] |
| M4 | 24,984 (Kuḍubi) | 33,000 | 12,734 | 19,200[d] (India) |
| R8 | 42,089 | | 32,783 | 43,300[a] |

| | | | |
|---|---|---|---|
| | 25,287 (Khārvi & Kuḍubi)<br>10,186 (Khārvi & Kuḍubi) | | |
| R30 | 44,668 | 53,576 | |
| | 17,330 (RSB & Kuḍubi) | | |
| | 14,032 (Kuḍubi) | | |
| U1 | 38769 (Kuḍubi) | 31,955 | 38,070[b] |
| | 34578 (Kuḍubi) | | |
| U2 | 33,362 | 42,805 | |
| | 13,533 (NKS) | | |
| | 11,025 (GSB) | | |
| U7 | 31,762 | 18,052 | 21,083[b] |
| U7a | 26,040 (GSB) | 16,718 | 41,400[d] (India)<br>18,200[e] |
| U8 | 32,552 (CSB) | 43,034 | |
| K1a | 25,044 | 18,433 | 27,400[c] |
| K1a+150 | 13,227 (CSB & RSB) | | 18,000[c] |

Clusters are represented within parenthesis
*Mutation rates as per [3]
### Based on coding region mutation rate $1.26 \pm 0.08 \times 10^{-8}$
[a] [4]
[b] [5]
[c] [6]
[d] [7] based on HVS1 region only
[e] [8]

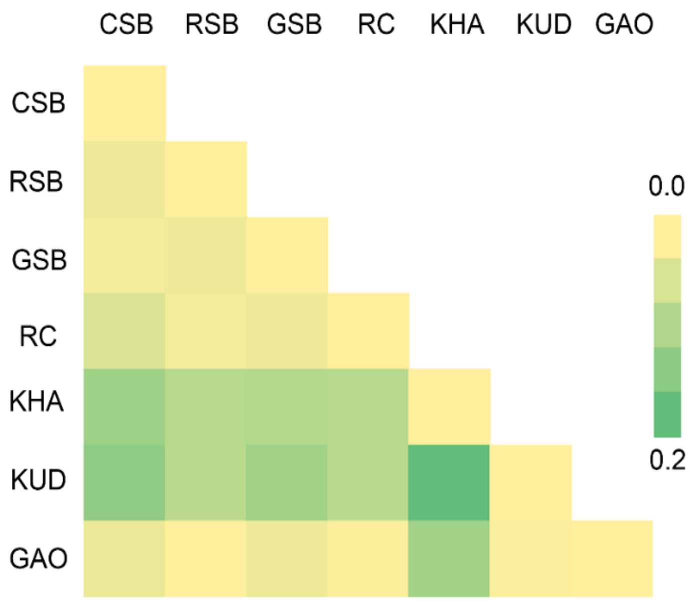


**Fig. S1: mtDNA Pairwise Fst matrix for Koṅkaṇī population**

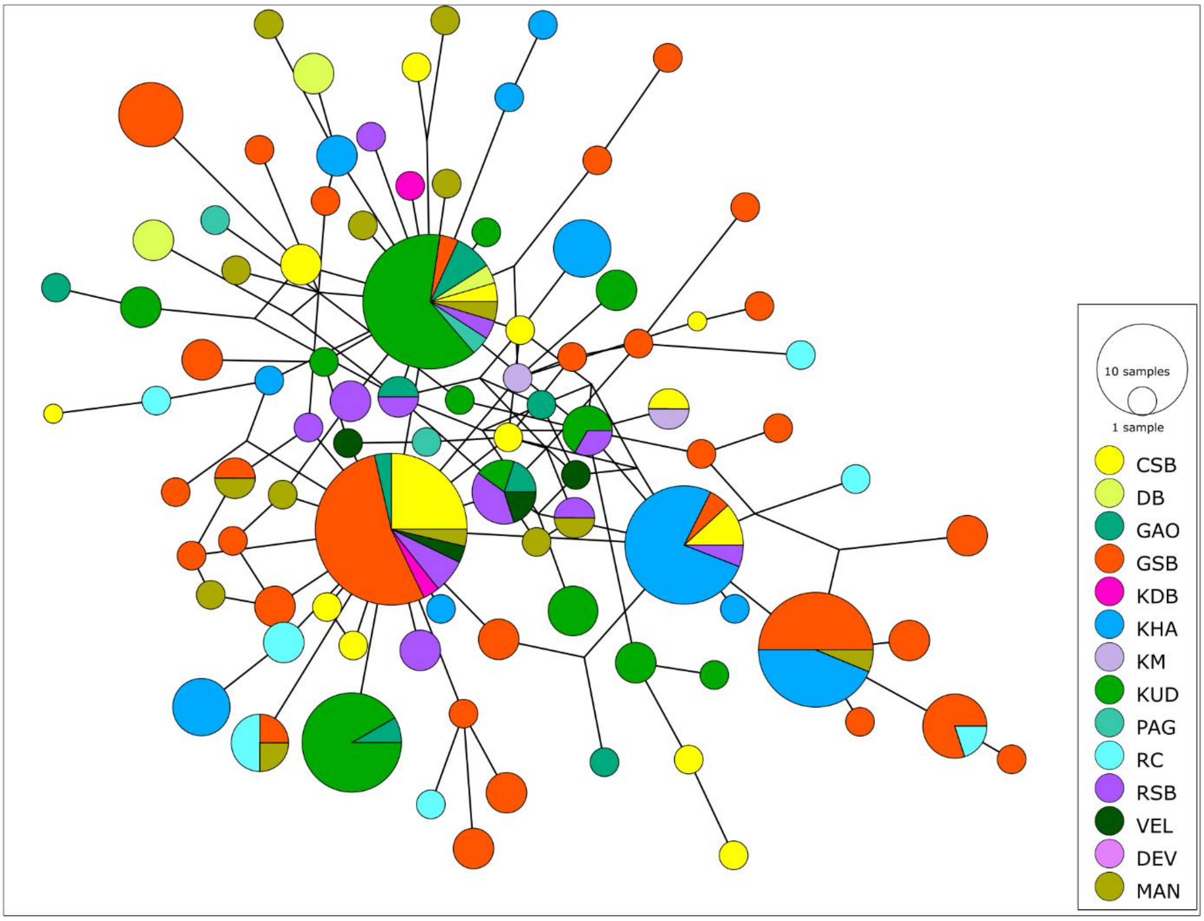


**Fig. S2: Haplotype network for Koṅkaṇī subpopulations**

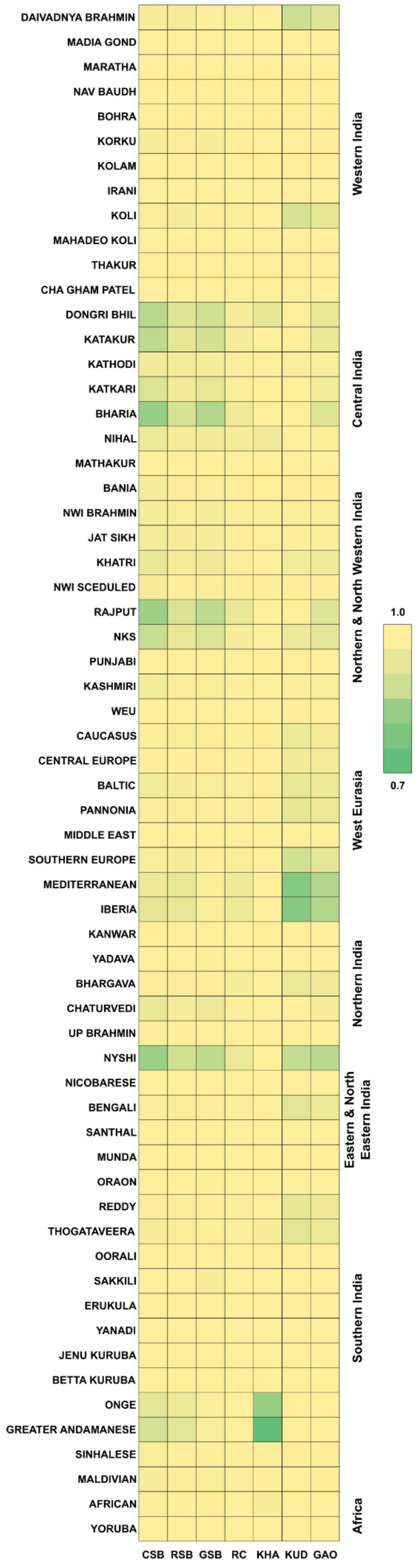


**Fig. S3 Heatmap showing average pairwise differences between Koṅkaṇī and other populations**

#### Bayesian Analysis

In order to estimate the time to the most recent common ancestor (TMRCA) for major haplogroups found in the studied population, we performed Bayesian analysis. We used complete mitogenome data obtained from 96 samples including all major haplogroups. Table 4 shows a comparison between our results and published estimates. Phylogenetic trees are shown in Fig.s S4-14.

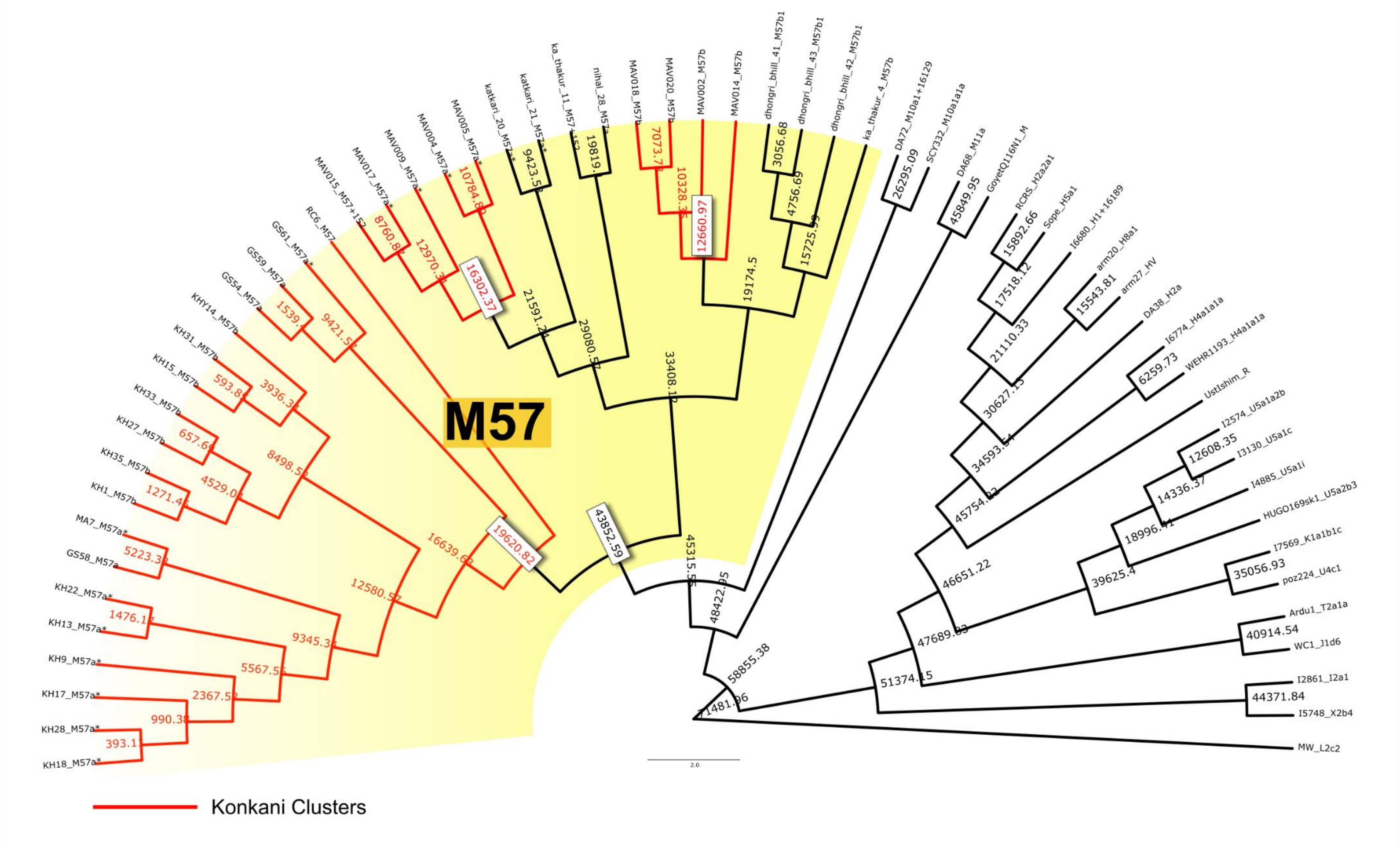

**Fig. S4: M57 phylogenetic tree**

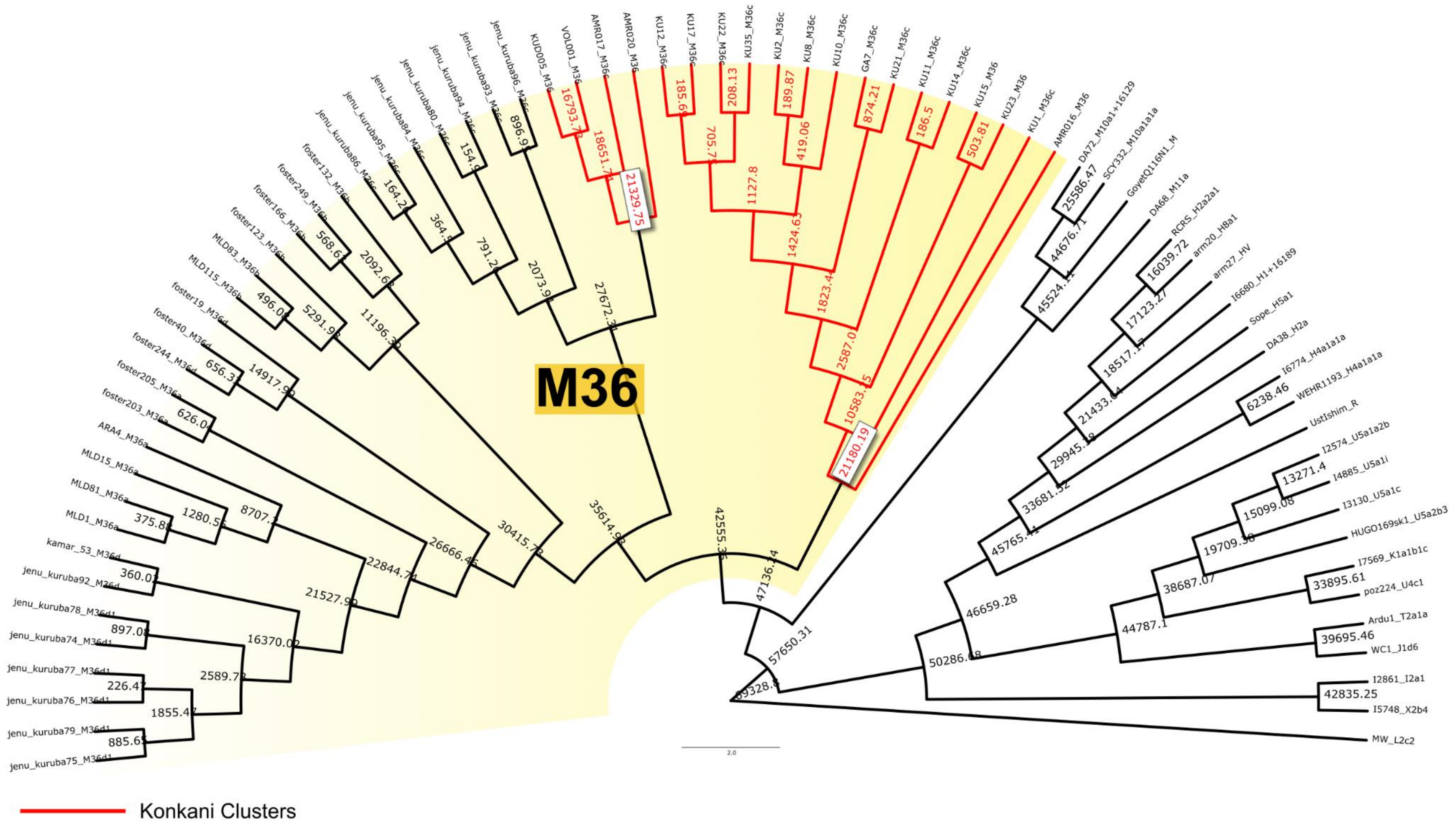


**Fig. S5: M36 phylogenetic tree**

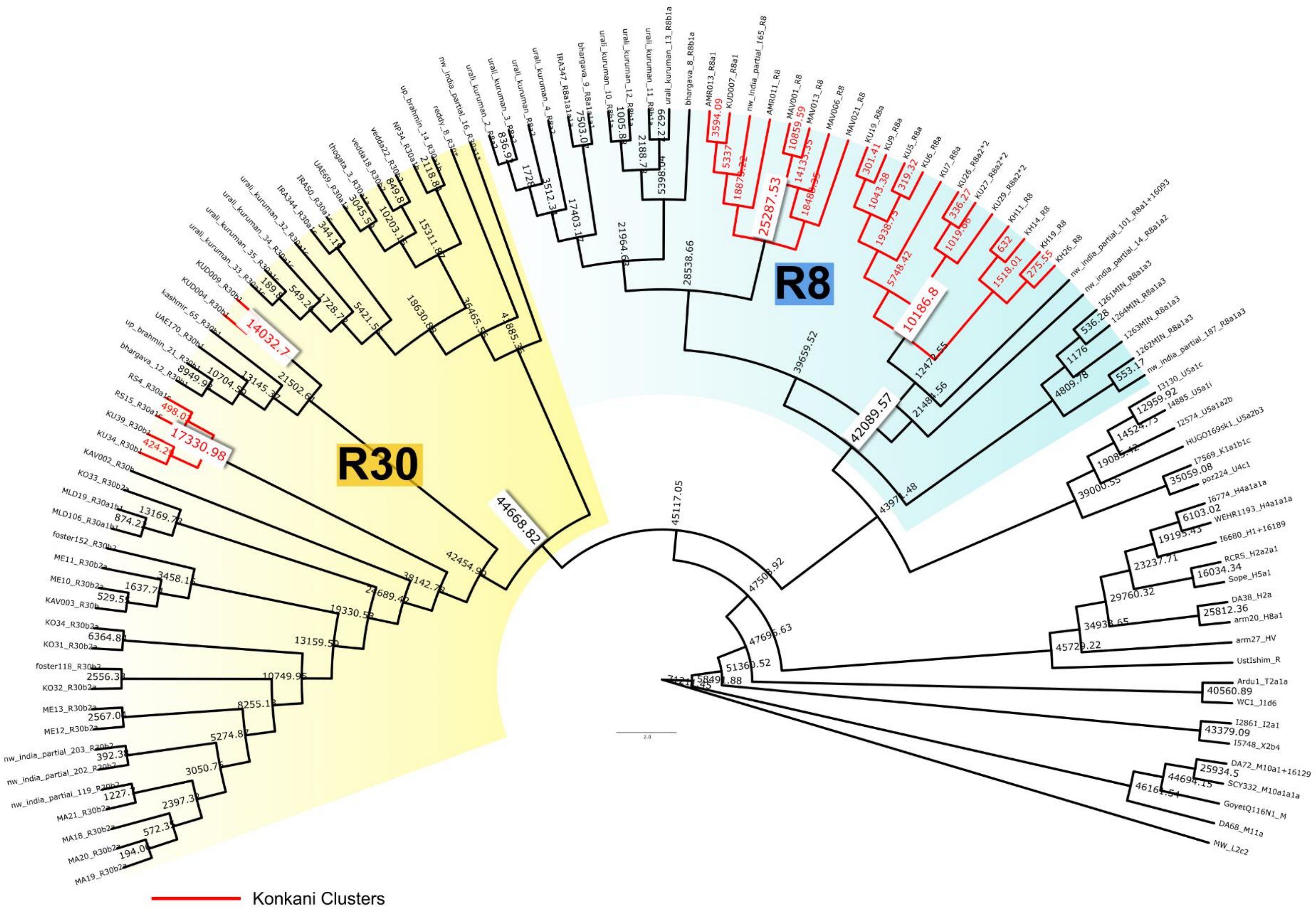


**Fig.S6: R8 and R30 phylogenetic tree**

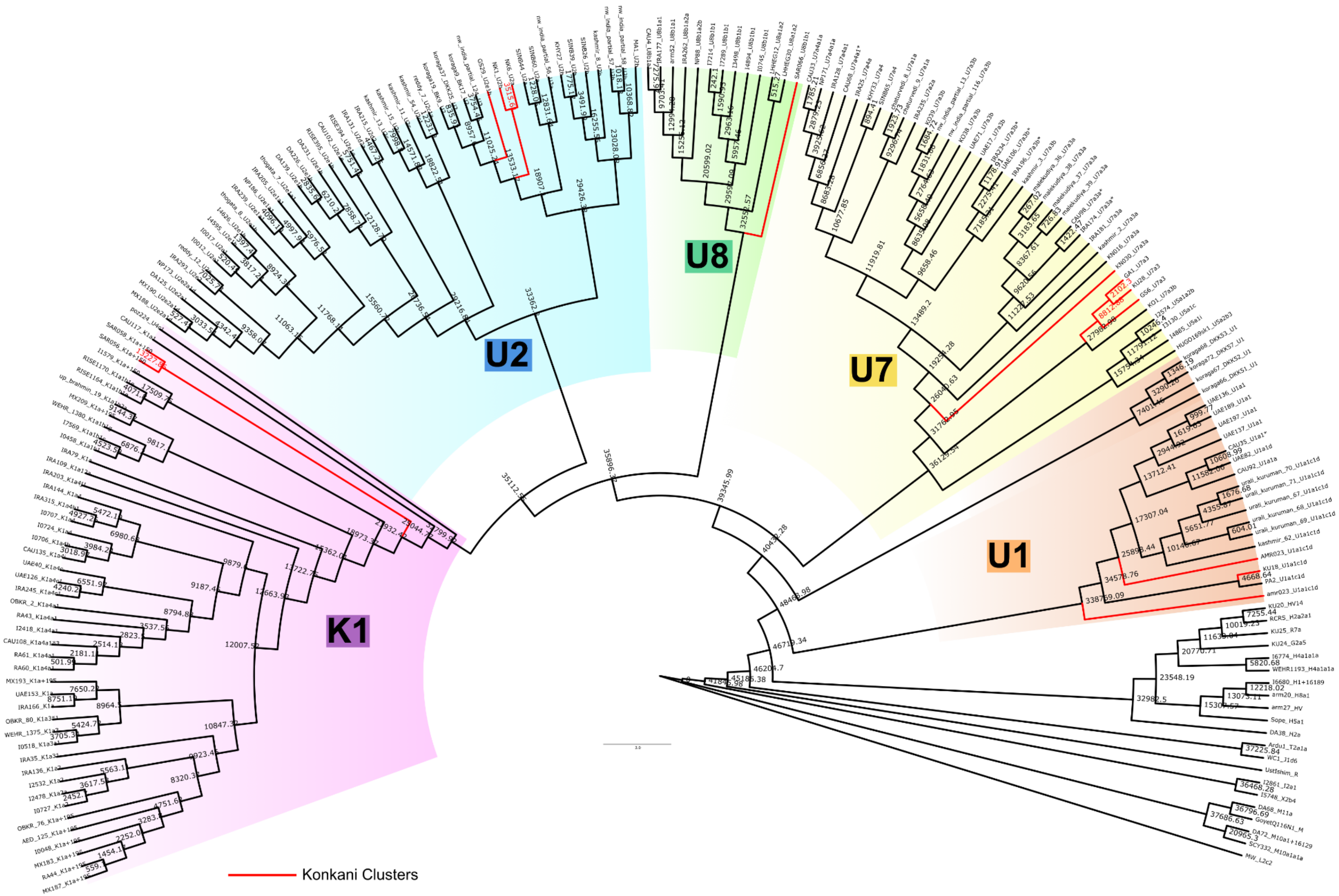


**Fig.S7: U and K1 phylogenetic tree**

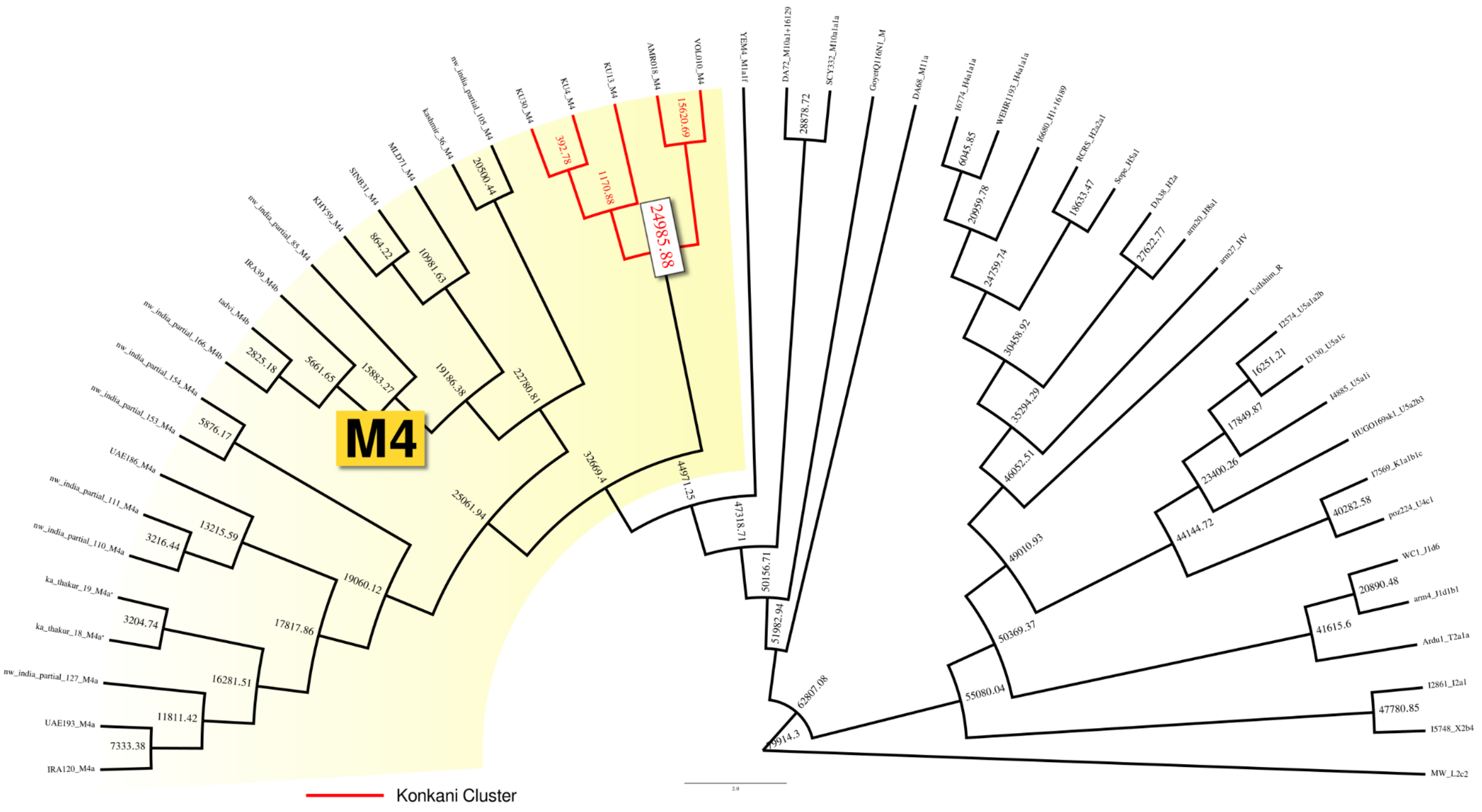


**Fig.S8: M4 Phylogenetic tree**

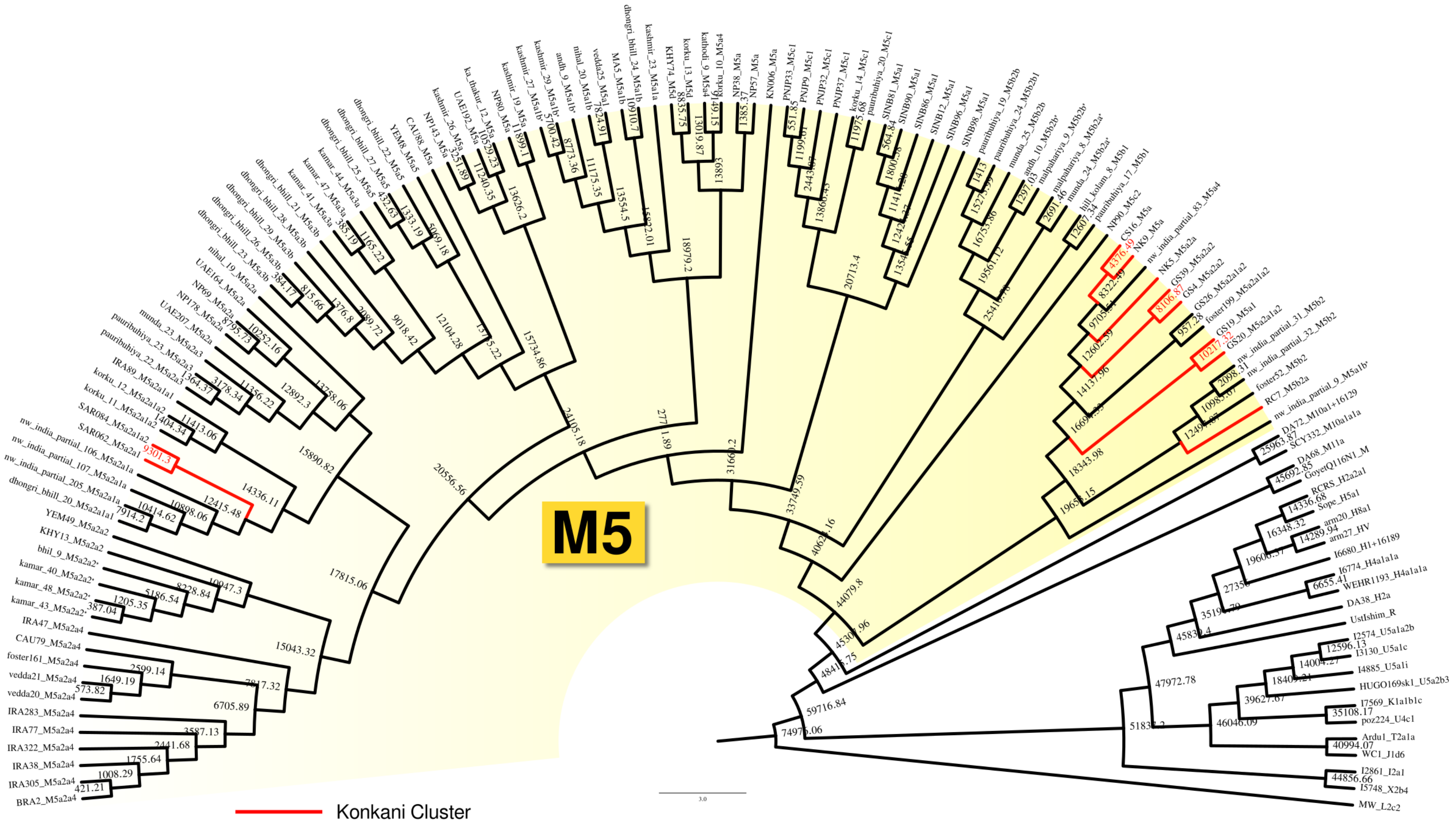


**Fig.S9: M5 Phylogenetic tree**

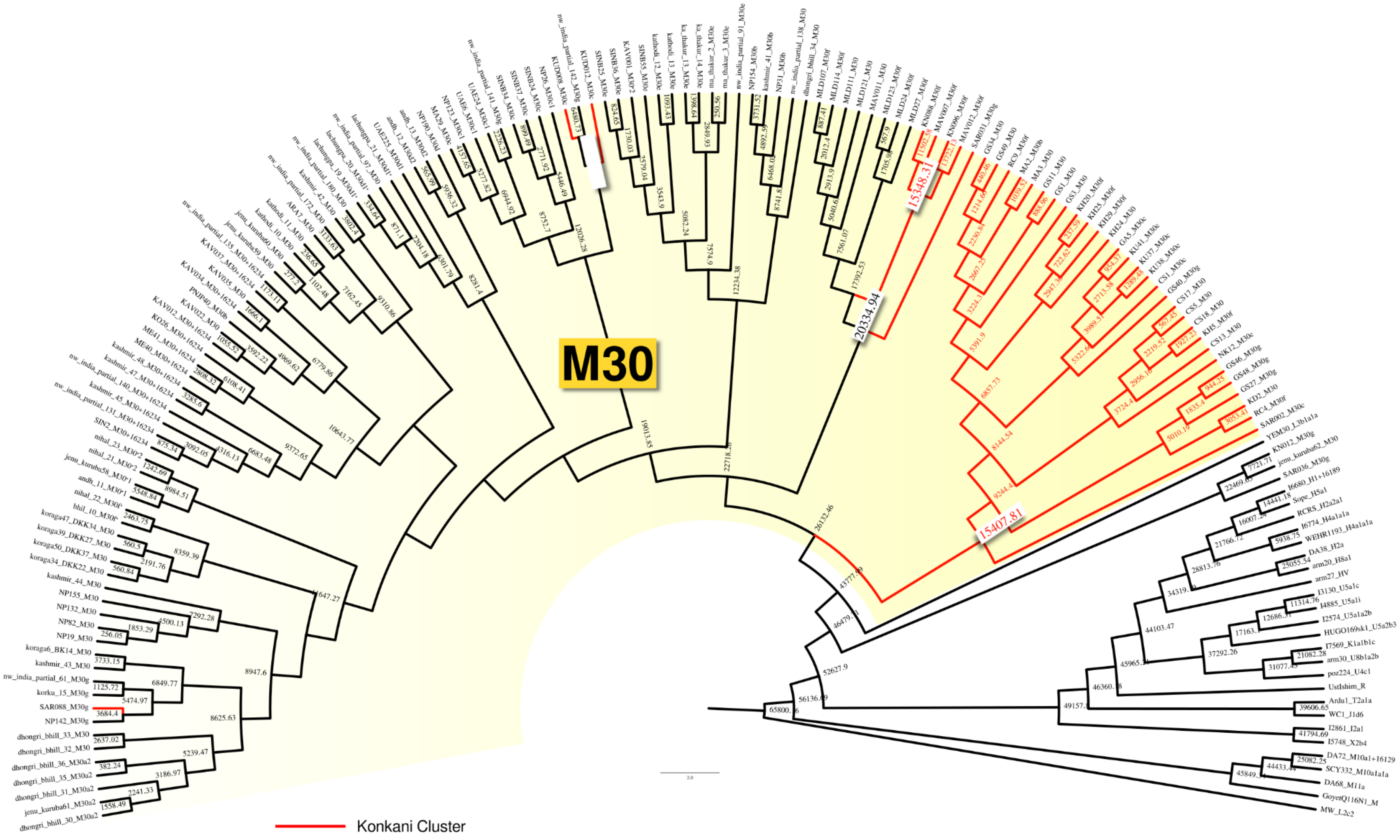


**Fig. S10: M30 Phylogenetic tree**

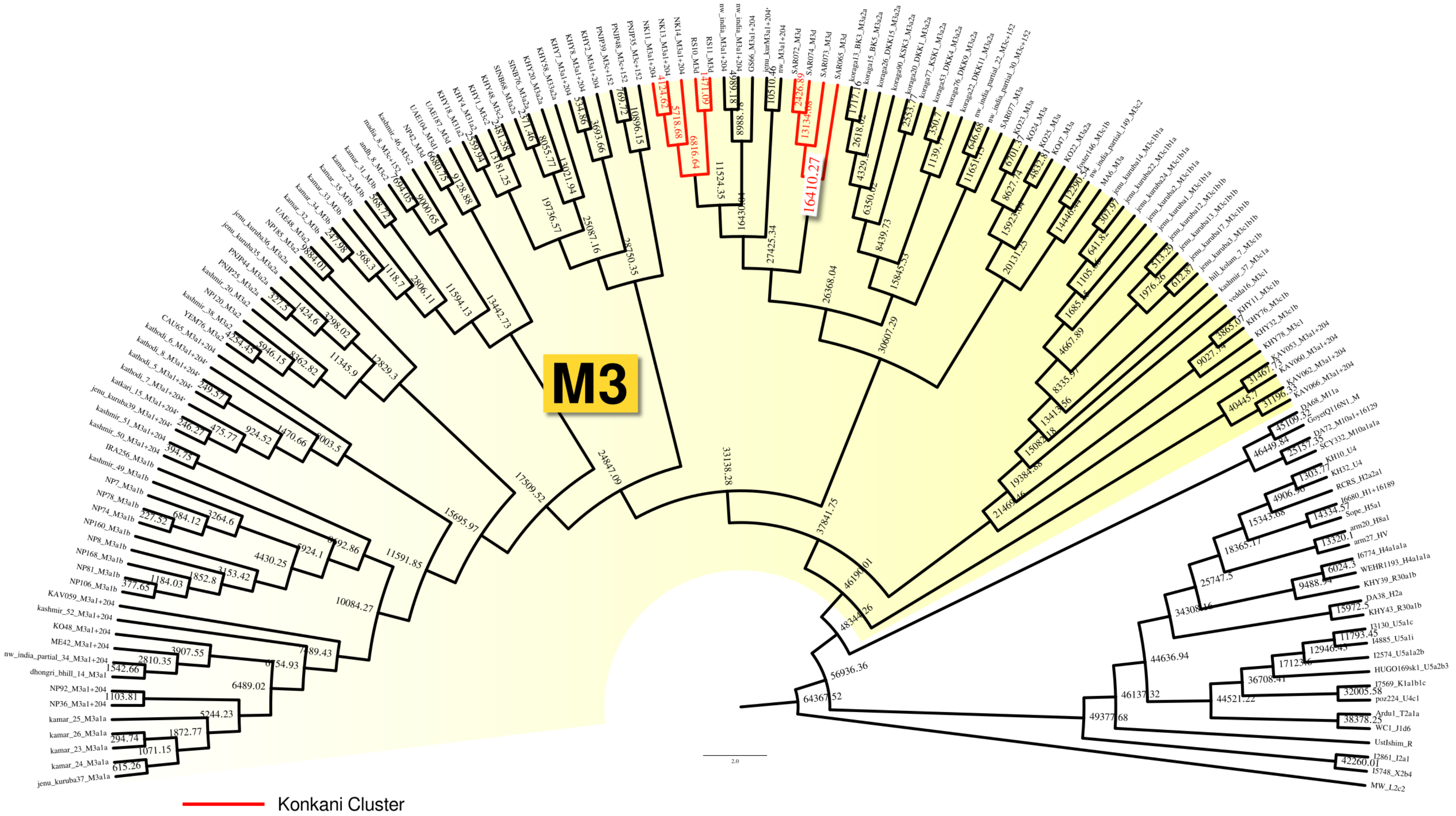


**Fig. S11: M3 Phylogenetic tree**

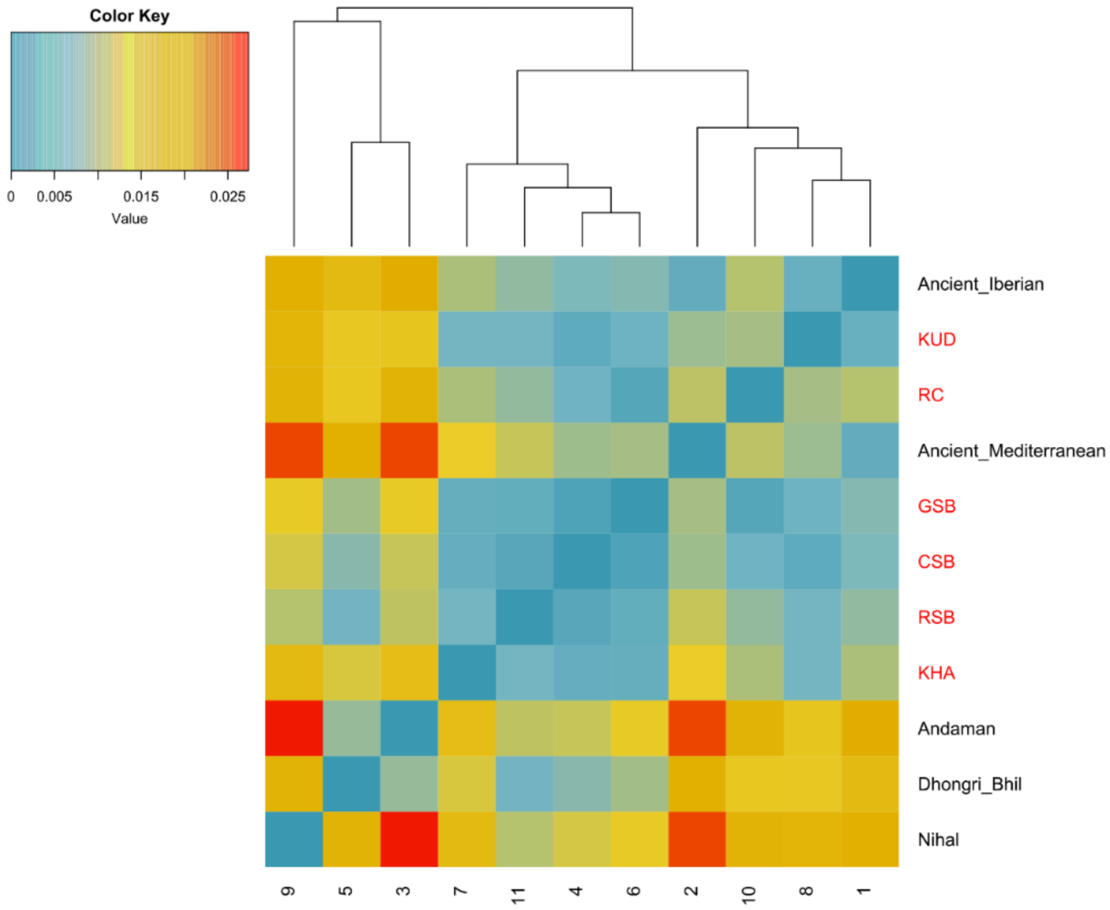


**Fig. S12 Heatmap of Nei's distance matrix based on the D-loop comparing Konkani subgroups with selected populations that showed similarities in haplogroup frequencies.**

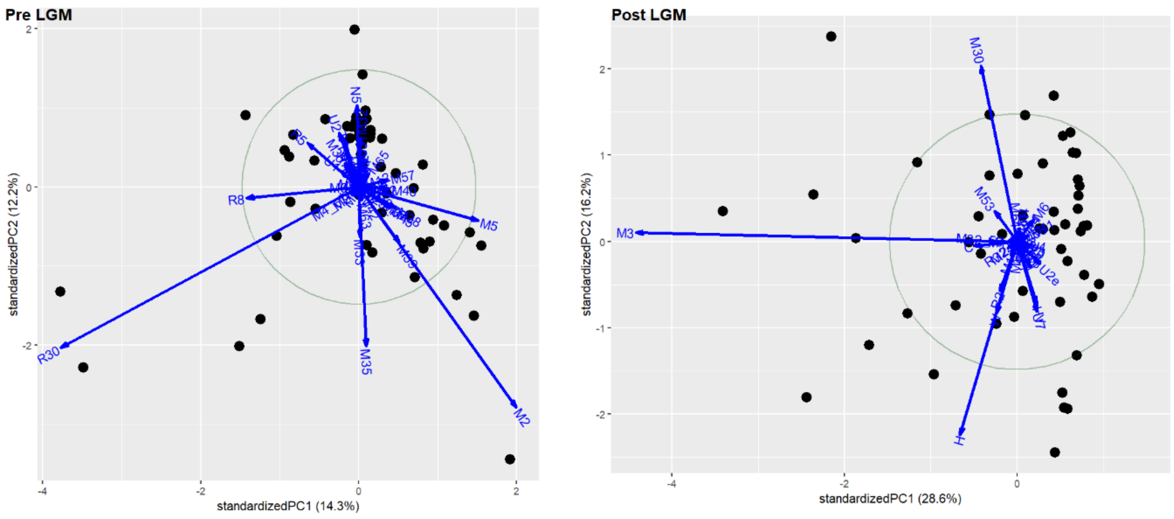


**Fig. S13 Biplots showing the haplogroup frequency contribution to the scatter in Pre and Post LGM era**
